# The UbiB family member Cqd1 forms a novel membrane contact site in mitochondria

**DOI:** 10.1101/2022.04.09.487722

**Authors:** Siavash Khosravi, Xenia Chelius, Ann-Katrin Unger, Johanna Frickel, Timo Sachsenheimer, Christian Lüchtenborg, Rico Schieweck, Britta Brügger, Benedikt Westermann, Till Klecker, Walter Neupert, Max E. Harner

## Abstract

Mitochondria are essential organelles of eukaryotic cells that are characterized by their unique and complex membrane system. They are confined from the cytosol by an envelope consisting of two membranes. Signals, metabolites, proteins and lipids have to be transferred across these membranes via proteinaceous contact sites to keep mitochondria functional. In the present study we identified a novel mitochondrial contact site that is formed by the inner membrane protein Cqd1 and the outer membrane proteins Por1 and Om14. Similar to the mitochondrial porin, Por1, Cqd1 is highly conserved, suggesting that this complex is conserved in form and function from yeast to human. Cqd1 is a member of the UbiB protein kinase-like family (also called aarF domain containing kinases). It was recently shown that Cqd1 in cooperation with Cqd2 controls the cellular distribution of coenzyme Q by a yet unknown mechanism. Our data suggest that Cqd1 in addition is involved in the homeostasis of phospholipids and contributes to the maintenance of mitochondrial morphology and architecture.

**Summary statement:** Here, we show that the conserved mitochondrial inner membrane protein Cqd1 interacts with the outer membrane proteins Por1 and Om14. Additionally, we provide evidence that Cqd1 is important for maintaining mitochondrial homeostasis.

## Introduction

Mitochondria are essential organelles in eukaryotic cells and are characterized by an envelope consisting of two membranes. In addition to their important role in the cellular energy metabolism, mitochondria perform a multitude of other functions, including the synthesis of proteins, iron-sulfur clusters and lipids. Specialized structures in the mitochondrial membranes, so-called contact sites, have to be present to enable mitochondria to perform all these functions. The term contact site often refers to inter-organellar contacts, for instance between mitochondria and the endoplasmic reticulum (ER) (Kornmann et al., 2009; Murley et al., 2015), mitochondria and vacuoles, the yeast equivalent of mammalian lysosomes (Gonzalez Montoro et al., 2018; John Peter et al., 2017), or mitochondria and lipid droplets (Pu et al., 2011). In addition, intra-mitochondrial contact sites between the mitochondrial inner and outer membrane can be observed (Tamura et al., 2019). Importantly, both kinds of contacts are crucial for mitochondrial functions (Khosravi and Harner, 2020; Klecker et al., 2014; Kornmann, 2020; Phillips and Voeltz, 2016; Renne and Hariri, 2021).

Most contacts between the mitochondrial inner and outer membrane depend on the mitochondrial contact site and cristae organizing system (MICOS), a multi-subunit protein complex in the mitochondrial inner membrane (Harner et al., 2011; Hoppins et al., 2011; von der Malsburg et al., 2011). This highly conserved complex interacts with at least six different proteins or complexes of the outer membrane: the Fzo1-Ugo1 complex (Harner et al., 2011), the Miro GTPases (Modi et al., 2019), Om45 (Hoppins et al., 2011), Por1 (Hoppins et al., 2011), the TOB (SAM) complex (Bohnert et al., 2012; Darshi et al., 2011; Harner et al., 2011; Korner et al., 2012; Xie et al., 2007; Zerbes et al., 2012) and the TOM complex (Bohnert et al., 2012; von der Malsburg et al., 2011; Zerbes et al., 2012).

In addition, several other intra-mitochondrial contact sites exist that do not depend on MICOS. Some of these have been known for decades, like the TIM23-TOM super complex (Chacinska et al., 2003; Dekker et al., 1997; Schleyer and Neupert, 1985; Schwaiger et al., 1987), or the mitochondrial fusion machineries (Fritz et al., 2001; Sesaki et al., 2003; Wong et al., 2003). Others were identified just recently, for instance the contact site formed by Por1 (yeast porin) and Mdm31 that was suggested to be important for the biosynthesis of cardiolipin (Miyata et al., 2018). Apparently, the functions of these intra-mitochondrial contacts are quite diverse, varying from protein import to lipid transport and the formation of mitochondrial architecture and morphology.

Members of the poorly characterized UbiB protein kinase-like family (aarF domain containing kinases) were recently implicated in mitochondrial membrane homeostasis (Awad et al., 2020; Johnson et al., 2005; Kemmerer et al., 2021; Mollet et al., 2008; Odendall et al., 2019; Reidenbach et al., 2018; Stefely et al., 2015). UbiB family members are defined by the presence of a protein kinase-like domain (PKL) of unknown function. *Saccharomyces cerevisiae* has three family members which all reside in mitochondria: Coq8 (Abc1), Cqd1 (Mco76) and Cqd2 (Mcp2) (Do et al., 2001; Morgenstern et al., 2017; Tan et al., 2013). The bacterial proteins UbiB *(Escherichia coli)* and AarF *(Providencia stuartii)* are essential for the synthesis of coenzyme Q (ubiquinone) (Macinga et al., 1998; Poon et al., 2000). Likewise, Coq8 and its mammalian homolog ADCK3 are also part of the coenzyme Q biosynthesis pathway (Do et al., 2001; Johnson et al., 2005; Mollet et al., 2008; Reidenbach et al., 2018; Stefely et al., 2015). While our work was in progress, an elegant study linked also Cqd1 and Cqd2 to coenzyme Q homeostasis. Deletion of *CQD1* results in excess export of coenzyme Q to extra-mitochondrial membranes, whereas deletion of *CQD2* leads to an accumulation of excess coenzyme Q in mitochondria. Thus, Cqd1 and Cqd2 apparently perform antagonistic roles in the distribution of coenzyme Q between mitochondria and other organelles (Kemmerer et al., 2021).

In the present study, we set out to functionally analyze the conserved mitochondrial protein encoded by the ORF *YPL109C,* which is now named Cqd1. We identified a MICOS-independent contact site formed by Cqd1 in the mitochondrial inner membrane and Por1 and Om14 in the outer membrane. Our characterization of the Δ*cqd1* deletion mutant suggests that Cqd1 contributes to phospholipid homeostasis in addition to its role in the regulation of the distribution of coenzyme Q. Moreover, we obtained evidence that defined levels of Cqd1 are important for mitochondrial morphology and architecture.

## Results

### Cqd1 is a mitochondrial inner membrane protein facing the intermembrane space

We first asked where Cqd1 is precisely located within mitochondria. *In silico* analysis using MitoProt II (Claros and Vincens, 1996) revealed that Cqd1 contains a potential N-terminal mitochondrial targeting sequence (MTS; amino acids 1-15) (Fig. 1A) (SACS MEMSAT2 Transmembrane Prediction (Jones et al., 1994)). In addition, a predicted transmembrane segment (amino acids 125-141) can be identified (Fig. 1A). We used an epitope-tagged construct (Cqd1-3xHA) to analyze the topology of Cqd1. Proteinase K (PK) accessibility and alkaline extraction assays using isolated mitochondria showed that it is behaving like Tim50 which is anchored in the inner membrane with an α-helical transmembrane domain (Fig. 1B and C). PK did not degrade Cqd1-3xHA in intact mitochondria, but the signal was largely gone when PK was added to hypotonically swollen mitochondria with a disrupted outer membrane. No shorter fragment appeared upon addition of PK to swollen mitochondria indicating that the C-terminal tag was degraded (Fig. 1B). Furthermore, Cqd1-3xHA remained in the pellet fraction upon alkaline extraction like the membrane proteins Tom70 and Tim50 (Fig. 1C). It should be noted that in Western blot analyses Cqd1-3xHA showed an apparent size of 66 kDa, instead of its calculated molecular weight of 76 kDa (untagged Cqd1 full length) or 74 kDa (untagged Cqd1 minus MTS) (Fig. 1B and C). We conclude that Cqd1 is present in the mitochondrial inner membrane with its major C-terminal part facing the intermembrane space.

**Figure 1.**
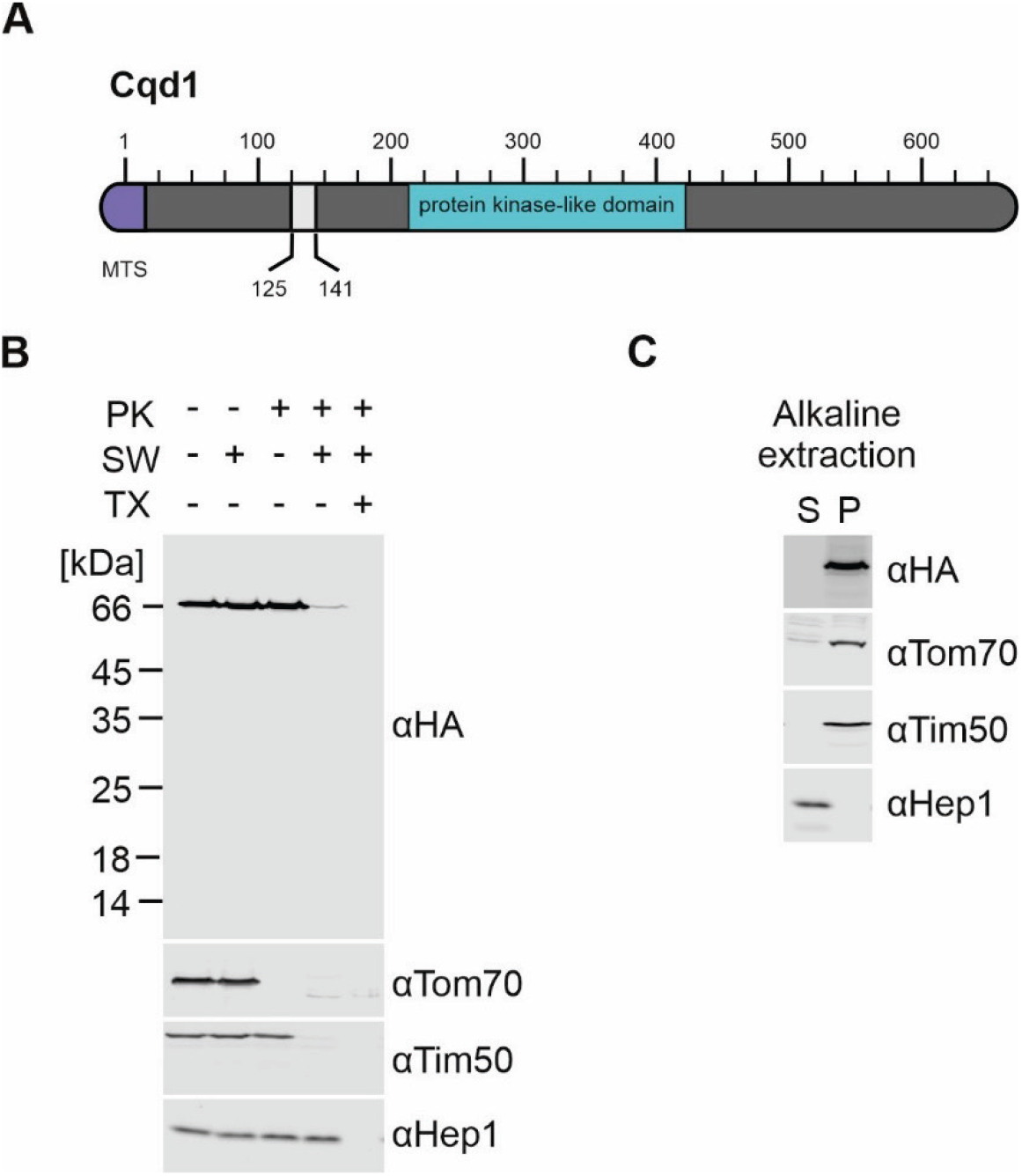
Cqd1 is a mitochondrial inner membrane protein exposing its C-terminus to the intermembrane space. **(A)** Schematic representation of Cqd1. Violet, mitochondrial targeting sequence (MTS; amino acids 1-15); light grey, predicted transmembrane domain (amino acids 125-141); turquoise, conserved protein kinase-like domain (amino acids 213-421). **(B)** Cqd1 exposes its C-terminus to the intermembrane space. Mitochondria isolated from a Cqd1-3xHA expressing strain were treated with isotonic buffer, subjected to swelling by incubation in hypotonic buffer to disrupt the outer membrane (SW), or lysed using a Triton X-100 containing buffer (TX). Proteinase K (PK) was added as indicated. Samples were analyzed by SDS-PAGE and immunoblotting. Tom70 was used as a marker for the outer membrane, Tim50 for the inner membrane and Hep1 for the matrix. **(C)** Cqd1 is an integral membrane protein. Isolated mitochondria were subjected to alkaline extraction to separate soluble and membrane proteins. Soluble proteins present in the supernatant (S) and membrane proteins in the pellet (P) were analyzed by SDS-PAGE and immunoblotting.

### The *CQD1* gene negatively interacts with *UPS1* and *CRD1*

Several high-throughput screens showed that cells lacking Cqd1 are able to grow on non-fermentable carbon sources (Qian et al., 2012; Stenger et al., 2020). Also, Kemmerer et al. (2021) observed growth of Δ*cqd1* cells under respiratory conditions in liquid media, albeit growth was reduced when coenzyme Q precursors were depleted. We generated a deletion mutant lacking Cqd1 and analyzed its growth phenotype under several conditions on agar plates. We observed that the Δ*cqd1* mutant grows like wild type on fermentable and non-fermentable carbon sources at normal and elevated temperatures (Fig. 2A) confirming that Cqd1 is not essential for respiratory growth.

**Figure 2.**
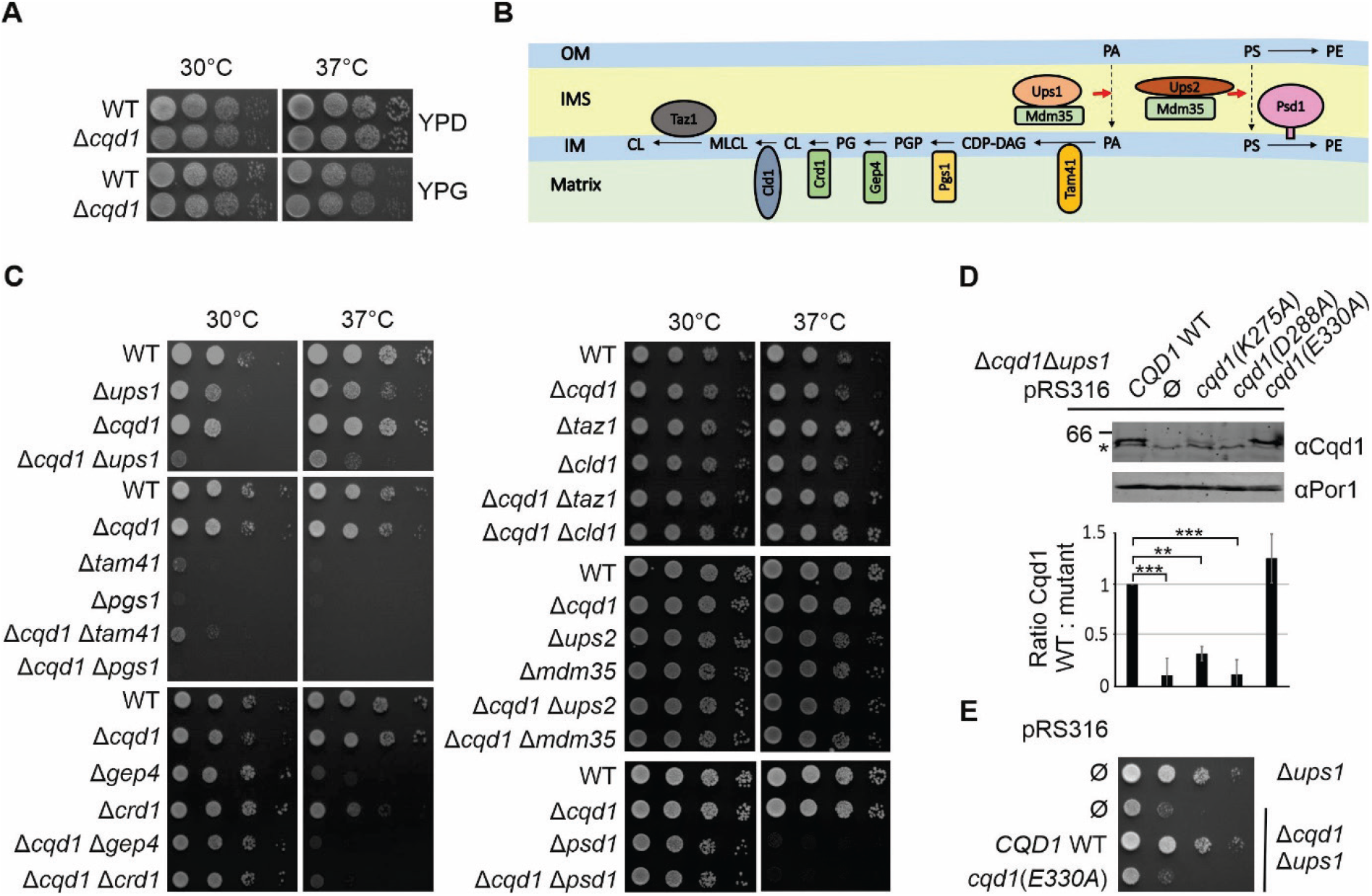
*CQD1* is a negative genetic interactor of *UPS1* and *CRD1*. **(A)** Deletion of *CQD1* does not result in a growth defect. Cells of wild type and a Δ*cqd1* deletion strain were grown to logarithmic growth phase in rich medium containing glucose as carbon source (YPD). Cell growth was analyzed by drop dilution assay on plates containing rich medium supplemented with either glucose (YPD) or glycerol (YPG) at 30°C or 37°C. **(B)** Schematic illustration of the mitochondrial phospholipid metabolism. OM, outer membrane; IMS, intermembrane space; IM, inner membrane; PA, phosphatidic acid; CDP-DAG, cytidine diphosphate diacylglycerol; PGP, phosphatidylglycerol phosphate; PG, phosphatidylglycerol; CL, cardiolipin; MLCL, monolysocardiolipin; PS, phosphatidylserine; PE, phosphatidylethanolamine. **(C)** Deletion of *CQD1* from cells lacking Ups1 or Crd1 results in a synthetic growth defect. Cells were treated as in (A) with the difference that they were shifted to synthetic medium containing glucose (SCD) 30 h before growth analysis on SCD plates. **(D)** The conserved amino acids K275 and D288 in the predicted protein kinase-like domain are important for the stability of Cqd1. Cells bearing either the empty plasmid (Ø) or plasmids carrying the respective *cqd1* alleles were grown in SCGal. Crude mitochondria were isolated and the Cqd1 levels were analyzed by immunoblotting using an anti-Cqd1 antibody. (Upper panel) Immunodecorations of one representative experiment. The asterisk indicates a cross reaction of the anti-Cqd1 antibody. (Lower panel) Quantitative analysis of Cqd1 steady state level in the different strains analyzed in three independent experiments. Quantification was done using Image Studio software. Error bars indicate standard deviation. Asterisks represent p-values calculated by one-way ANOVA with subsequent Tukey’s multiple comparison test (**p≤0.01; ***p≤0.001). **(E)** Glutamic acid 330 is essential for the function of Cqd1. Cells bearing either the empty plasmid or a plasmid carrying *CQD1* wild type (WT) or *cqd1(E330A)* alleles were treated as in (C).

Mitochondria are intensively involved in the cellular phospholipid metabolism as they generate cardiolipin and phosphatidylethanolamine. Interestingly, two independent high-throughput studies revealed a genetic interaction of *CQD1* with *UPS1* (Costanzo et al., 2016; Hoppins et al., 2011). Ups1 is responsible for transport of phosphatidic acid across the intermembrane space (Fig. 2B) (Connerth et al., 2012; Tamura et al., 2012). Thus, we generated double deletion mutants lacking *CQD1* and genes encoding proteins involved in mitochondrial phospholipid metabolism, including *CLD1, CRD1, GEP4, MDM35, PGS1, PSD1, TAM41, TAZ1, UPS1* and *UPS2* (Fig. 2B). Growth analysis revealed a strong defect of the double deletion mutant Δ*cqd1* Δ*ups1* specifically on synthetic medium (Fig. 2C). A weaker yet reproducible growth defect at 37°C could be detected for the double deletion mutant Δ*cqd1* Δ*crd1.* All other double deletion mutants did not show an enhanced growth phenotype. The growth defect of the single deletion mutants Δ*tam41* and Δ*pgs1* was so strong that we cannot judge whether the additional deletion of *CQD1* further reduces cell growth (Fig. 2C).

Cqd1 is a member of the highly conserved UbiB family (Kemmerer et al., 2021; Stefely et al., 2015) and thus shows a predicted protein kinase-like domain (amino acids 213-421, Saccharomyces Genome Database, SGD (Cherry et al., 2012)) (Fig. 1A). Therefore, we asked whether the protein kinase-like domain of Cqd1 is important for its function. We generated mutants in conserved amino acid residues within this domain (K275A, D288A and E330A) to inhibit ATP binding similar to what was done before in studies on Cqd2 and Coq8 (Odendall et al., 2019; Reidenbach et al., 2018; Stefely et al., 2015). Of note, introduction of the point mutations K275A or D288A led to strongly reduced steady state levels of Cqd1 indicating reduced expression or stability of the protein (Fig. 2D). Therefore, we did not further analyze these alleles. In contrast, the introduction of the point mutation E330A did not affect the steady state level of Cqd1 (Fig. 2D). While expression of wild type Cqd1 largely restored the growth of the double deletion mutant Δ*cqd1* Δ*ups1,* the *cqd1(E330A)* allele did not, indicating that ATP binding of Cqd1 is important for its function (Fig. 2E). These results are in accordance with observations recently made by Kemmerer et al. (2021) who reported that alleles carrying mutations of conserved residues of the kinase-like domain were unable to rescue Δ*cqd1* growth defects.

In sum, we could manually confirm the negative genetic interaction of *CQD1* and *UPS1* and identified *CRD1* as a novel negative interactor of *CQD1.* Interestingly, both genetic interactors are involved in biosynthesis of the phospholipid cardiolipin. Furthermore, Cqd1 activity depends on an intact protein kinase-like domain.

### Cqd1 contributes to mitochondrial lipid homeostasis

Next, we analyzed the phospholipid composition of mitochondria isolated from wild type yeast, the Δ*ups1* or Δ*cqd1* single deletion strains and the Δ*cqd1* Δ*ups1* double deletion mutant by mass spectrometry. Consistent with previous results (Connerth et al., 2012), deletion of *UPS1* resulted in a severe reduction of cardiolipin and monolysocardiolipin (Fig. 3, Suppl. Table 1). Also, we observed a significant reduction of phosphatidic acid in Δ*ups1*, which might depend on growth conditions (Eiyama et al., 2021). Interestingly, the lipidomics analysis of mitochondria obtained from Δ*cqd1* revealed a significant reduction of phosphatidic acid while other phospholipids remained unchanged (Fig. 3, Suppl. Table 1). The double deletion mutant Δ*cqd1* Δ*ups1* did not show a synthetic phenotype. We suggest that Cqd1 is involved not only in maintenance of mitochondrial coenzyme Q levels (Kemmerer et al., 2021), but also modulates the levels of phosphatidic acid (Fig. 3, Suppl. Table 1).

**Figure 3.**
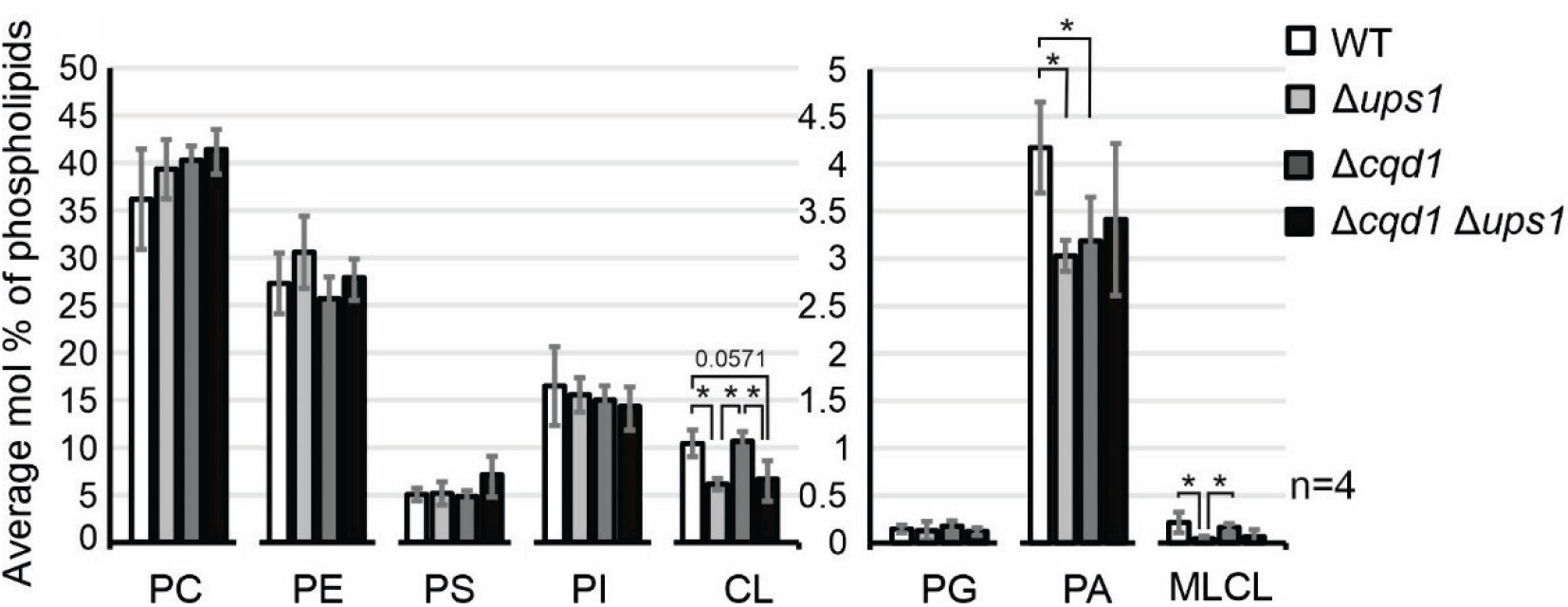
Cqd1 is involved in mitochondrial phospholipid homeostasis. Strains were grown in synthetic medium containing glycerol (SCG) and mitochondria were purified by sucrose gradient centrifugation. Phospholipids were extracted and analyzed by mass spectrometry. The level of each phospholipid species (% of total phospholipids) is shown as a mean of four independently performed experiments with standard deviation bars. Asterisks represent p-values obtained by Mann Whitney test (*p≤0.05). PC, phosphatidylcholine; PE, phosphatidylethanolamine; PS, phosphatidylserine; PI, phosphatidylinositol; CL, cardiolipin; PG, phosphatidylglycerol; PA, phosphatidic acid; MLCL, monolysocardiolipin.

### Mitochondrial biogenesis and dynamics are impaired in the Δ*cqd1* Δ*ups1* double mutant

The drastic growth phenotype of the Δ*cqd1* Δ*ups1* double mutant suggests that this strain struggles with severe problems in essential mitochondrial functions which might be caused by a lack of mitochondrial proteins or complexes involved in mitochondrial dynamics, architecture, protein import and/or respiration. Therefore, we analyzed the steady state levels of various proteins involved in these processes, including Fzo1, Ugo1, Mic27, Mic60, Tim23, Tim50, Rip1 and Cor2. Also, we tested assembly of the TOM complex and the respiratory chain super complexes by native gel electrophoresis. However, a comparison of the Δ*cqd1* Δ*ups1* double mutant with wild type and the single deletion mutants revealed no significant defects (Fig. 4A and B).

**Figure 4.**
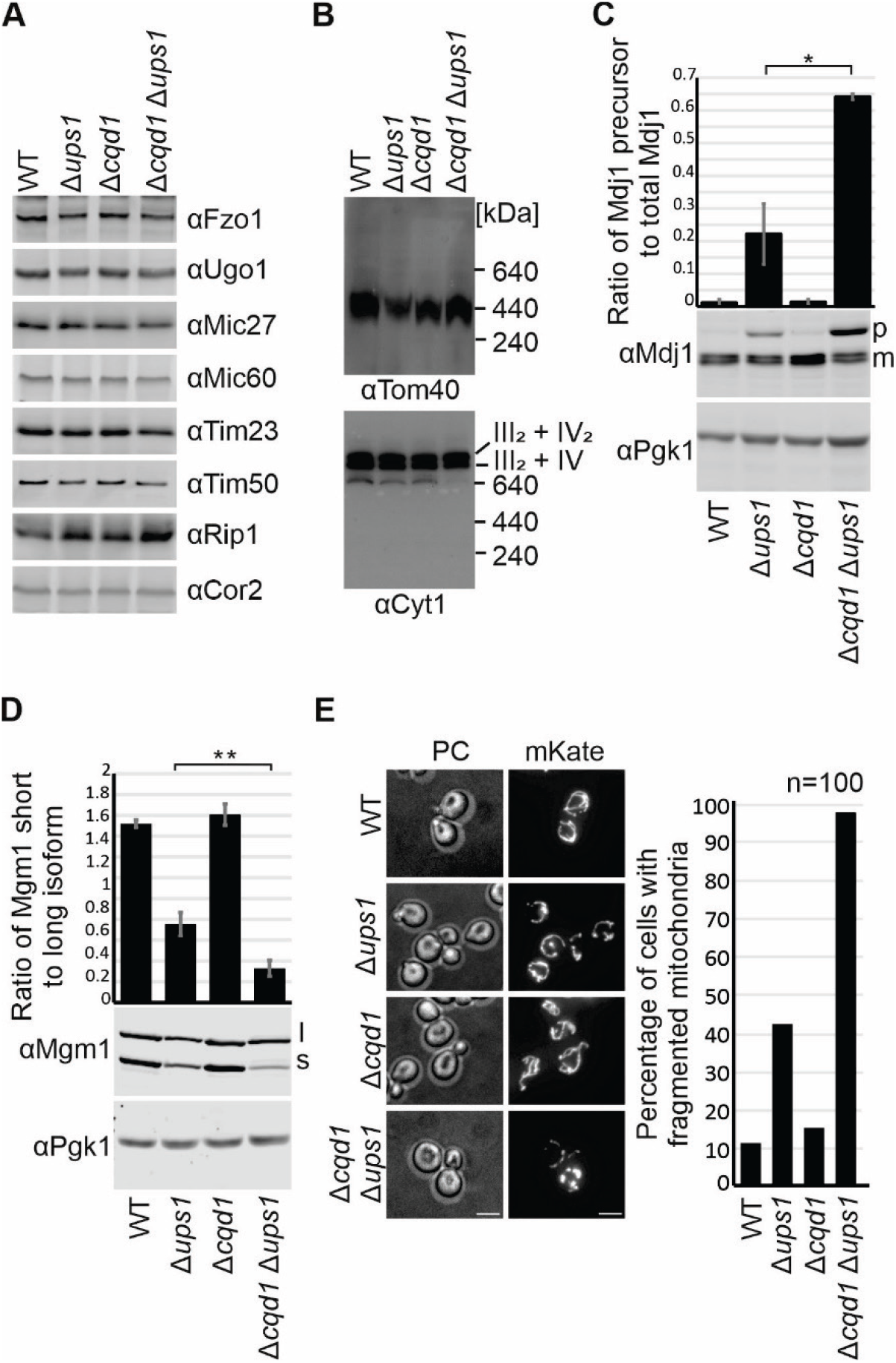
Simultaneous deletion of *CQD1* and *UPS1* impairs mitochondrial protein import and dynamics. **(A)** Analysis of steady state levels of mitochondrial proteins in wild type cells and cells lacking Ups1, Cqd1, or both. Cells were grown in synthetic complete medium containing galactose (SCGal), and whole cell extracts were analyzed by immunoblotting. **(B)** Formation of mitochondrial protein complexes. Strains were grown in SCG. Isolated mitochondria were lysed in digitonin-containing buffer (3% w/v) and cleared lysates were subjected to BN-PAGE. The assembly of the TOM complex and respiratory chain super complexes were analyzed by immunoblotting using antibodies against Tom40 or Cyt1. **(C)** Deletion of *CQD1* in cells lacking Ups1 exacerbates accumulation of the precursor of Mdj1. Cells were grown in SCD, whole cell lysates were prepared and analyzed by immunoblotting with specific antibodies. Pgk1 served as a loading control. p, precursor of Mdj1; m, mature form of Mdj1. The quantification was obtained from three independent experiments and shows means of the ratio of the Mdj1 precursor to the total amount of Mdj1. Quantification was done using Image Studio software. Error bars indicate standard deviation. Asterisk represents p-value obtained by unpaired Student’s t test (*p≤0.05). **(D)** Simultaneous deletion of *CQD1* and *UPS1* leads to strongly reduced processing of Mgm1. Whole cell lysates from cells grown in SCD were analyzed by immunoblotting. l, long isoform of Mgm1; s, short isoform of Mgm1. The quantification was obtained from three independent experiments and shows means of the ratio of the short form to the long form of Mgm1. Quantification was done using Image Studio software. Asterisks represent p-values obtained by unpaired Student’s t test (**p≤0.01). **(E)** Mitochondria in cells lacking Ups1 and Cqd1 are highly fragmented. Mitochondria were labeled by expression of mKate targeted to mitochondria. Cells were grown in YPD and shifted to SCD, harvested in their logarithmic growth phase and immobilized on slides covered with concanavalin A. Scale bars, 4 μm. PC, phase contrast. For quantification 100 cells were counted for each strain.

Previous studies showed that deletion of *UPS1* compromises two processes that both depend on the mitochondrial membrane potential. First, the import of the mitochondrial matrix-localized chaperone Mdj1 is impaired resulting in the accumulation of unprocessed precursor (Tamura et al., 2009). And second, processing of Mgm1, a large inner membrane-associated GTPase required for mitochondrial fusion, is reduced resulting in the accumulation of the long isoform, l-Mgm1 (Sesaki et al., 2006). Interestingly, deletion of *CQD1* did neither affect the import of Mdj1 nor processing of Mgm1. However, we found that both defects observed in the Δ*ups1* single mutant are exacerbated in the Δ*cqd1* Δ*ups1* double mutant. We observed a strong accumulation of Mdj1 precursor protein and an almost complete inhibition of Mgm1 processing (Fig. 4C and D). Previous studies showed that a balanced ratio of processed s-Mgm1 to unprocessed l-Mgm1 is necessary for mitochondrial fusion (DeVay et al., 2009; Zick et al., 2009). Accordingly, we observed a wild type like mitochondrial network in the Δ*cqd1* single deletion mutant. In the Δ*cqd1* Δ*ups1* double deletion mutant, however, the mitochondrial network appeared almost completely fragmented, indicating impairment of mitochondrial fusion (Fig. 4E). Thus, the relative mild phenotypes of the Δ*ups1* single deletion mutant regarding cell growth, generation of membrane potential and morphology of mitochondria (Sesaki et al., 2006; Tamura et al., 2009) are enhanced by additional deletion of the *CQD1* gene.

### Cqd1 and Cqd2 have antagonistic roles in mitochondrial biogenesis

A previous study indicated opposing functions of Cqd1 and Cqd2 (Kemmerer et al., 2021). Therefore, we generated all possible single, double and triple deletion mutants to analyze the interaction network of *CQD1, CQD2* and *UPS1.* Strikingly, growth analysis of the triple deletion mutant Δ*cqd1* Δ*ups1* Δ*cqd2* revealed that deletion of *CQD2* in the Δ*cqd1* Δ*ups1* background completely rescued the growth phenotype (Fig. 5A). At the same time, the accumulation of Mdj1 precursor protein and the impaired Mgm1 processing were largely rescued (Fig. 5B and C). This suggests that the equilibrium of the putative kinases Cqd1 and Cqd2 is particularly important when mitochondrial membrane lipid composition is disturbed through deletion of *UPS1.*

**Figure 5.**
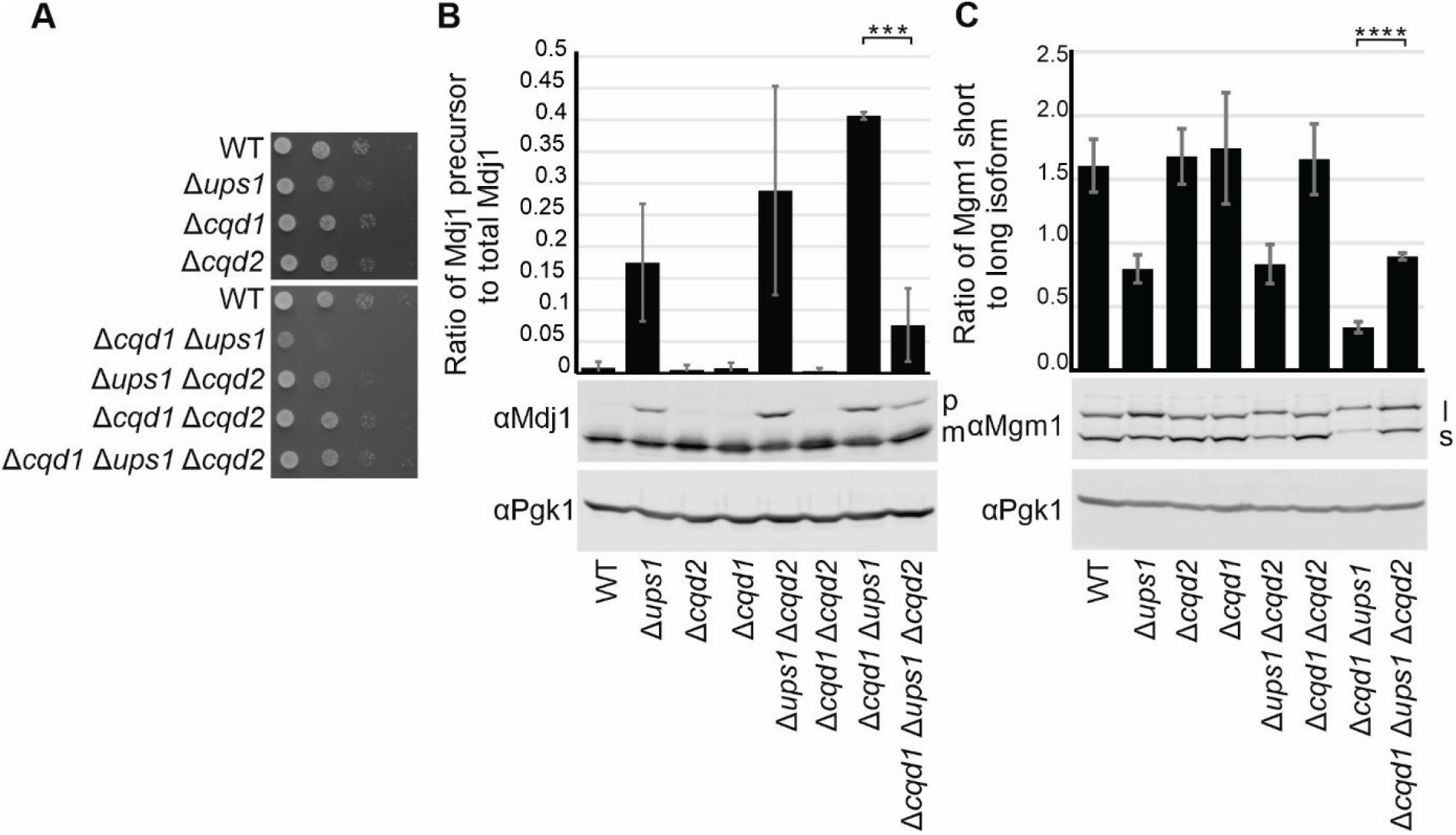
Deletion of *CQD2* restores the phenotypes of a Δ*cqd1* Δ*ups1* double deletion. **(A)** Deletion of *CQD2* in the Δ*cqd1* Δ*ups1* background rescues the growth phenotype. Strains were grown to logarithmic phase and growth was analyzed by drop dilution assay on SCD plates at 30°C. **(B+C)** Mdj1 import and Mgm1 processing is restored in the Δ*cqd1* Δ*ups1* Δ*cqd2* triple mutant. Cells were grown in SCD, whole cell lysates were prepared and analyzed by immunoblotting. Pgk1 served as a loading control. **(B)** p, precursor of Mdj1; m, mature form of Mdj1. The quantification (Image Studio software) was obtained from three independent experiments and shows means of the ratio of the Mdj1 precursor to the total amount of Mdj1. **(C)** l, long isoform of Mgm1; s, short isoform of Mgm1. The quantification (Image Studio software) was obtained from three independent experiments and shows means of the ratio of the short form to the long form of Mgm1. Error bars indicate standard deviation. Asterisks represent p-values obtained by unpaired Student’s t test (***p≤0.001; ****p≤0.0001).

### Cqd1 is part of a MICOS-independent contact site

Our results described above and the observations reported by Kemmerer et al. (2021) suggest that Cqd1 is involved in mitochondrial lipid homeostasis. We reasoned that a membrane contact site would be ideally suited for such a function. To test whether Cqd1 is a contact site protein, we used an established method to fractionate mitochondria, which is based on sonication, dounce homogenization and subsequent density gradient centrifugation of mitochondrial particles (Harner et al., 2011). Strikingly, Western blot analysis of mitochondrial fractions revealed that the distribution of Cqd1 is clearly distinguishable from both the outer membrane marker Tom40 and the inner membrane marker Tim17. Cqd1 was enriched in fractions of intermediate density very similar to the MICOS subunit Mic27 (Fig. 6A and Suppl. Fig. 1).

**Figure 6.**
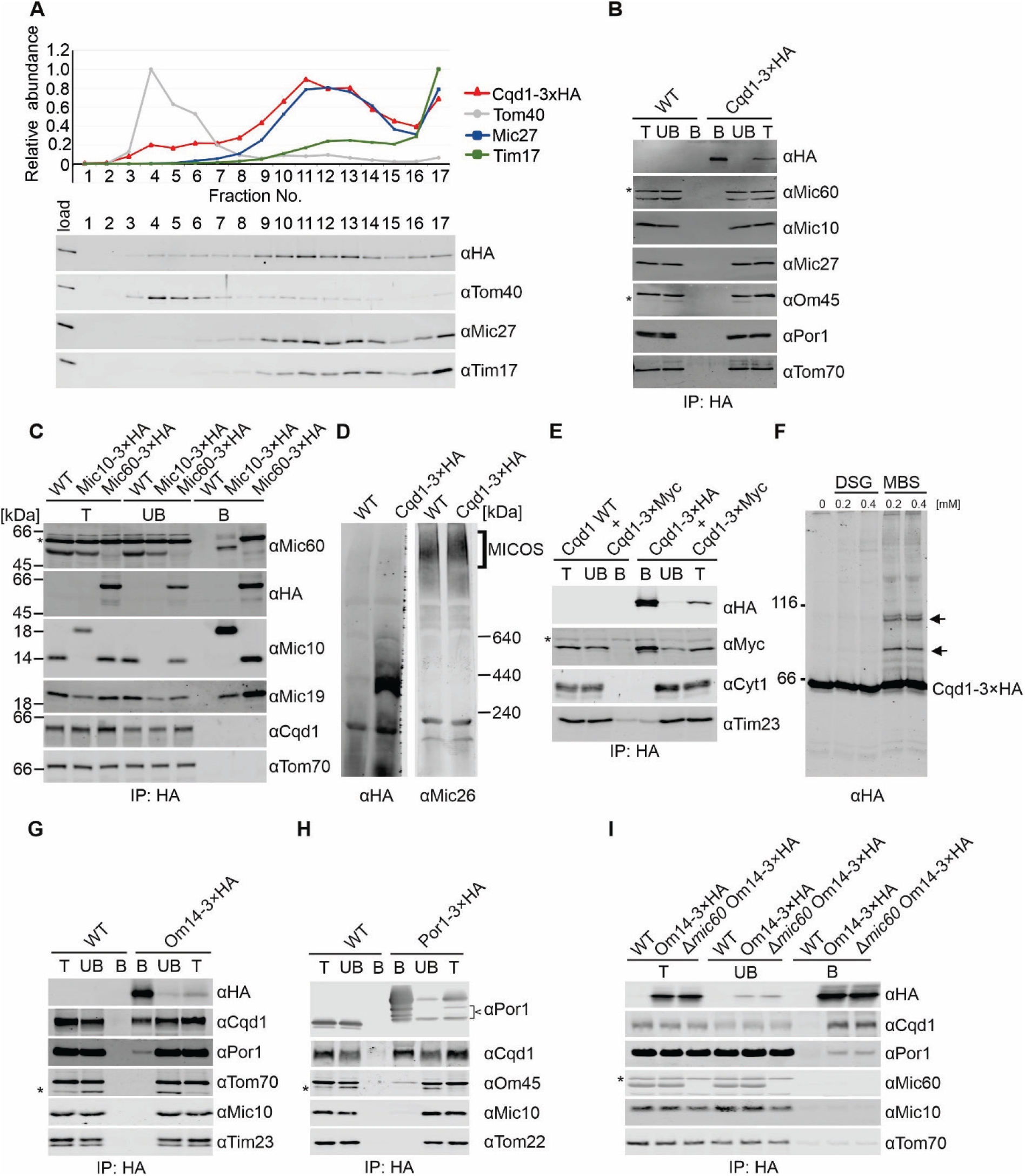
Cqd1 forms a novel contact site with Om14 and Por1. **(A)** Cqd1 is enriched in contact site fractions. Mitochondria of a Cqd1-3xHA expressing strain were isolated, subjected to osmotic shrinking, sonication, and sucrose density gradient centrifugation. The gradient was fractionated, proteins were subjected to TCA precipitation and analyzed by immunoblotting. (Top) The graph shows mean values of three independent experiments for the distribution of Cqd1-3xHA and the marker proteins for the outer membrane (Tom40), the inner membrane (Tim17) or contact sites (Mic27). Error bars indicating standard deviations are shown in Suppl. Fig. 1. (Bottom) Immunodecorations of one representative experiment. **(B)** Mic10 and Mic60 do not co-precipitate with Cqd1. Mitochondria of wild type and a yeast strain expressing Cqd1-3xHA were isolated and lysed in digitonin containing buffer (1% w/v). Lysates were subjected to immunoprecipitation using anti-HA affinity agarose. The indicated fractions were analyzed by SDS-PAGE and immunoblotting. T, total lysate (5%); UB, unbound protein (5%); B, bound protein (100%). Asterisks indicate cross reactions of the antibodies against Mic60 or Om45. **(C)** Cqd1 does not co-precipitate with Mic10 or Mic60. Mitochondria of wild type or yeast strains expressing Mic10-3xHA or Mic60-3xHA were analyzed as in (B). T, total lysate (2.5%); UB, unbound protein (2.5%); B, bound protein (100%). The asterisk indicates a cross reaction of the anti-Mic60 antibody. Since the cross reaction of the Mic60 antibody shows the same size as Mic60-3xHA, a decoration of this membrane fragment with an anti-HA antibody is presented additionally. **(D)** Cqd1 forms high molecular weight complexes. Mitochondria isolated from wild type and a yeast strain expressing Cqd1-3xHA were solubilized in digitonin (3% w/v). Cleared lysates were subjected to BN-PAGE. Cqd1-3xHA containing complexes were detected by immunoblotting with an anti-HA antibody. Analysis of the MICOS complex using an anti-Mic26 antibody served as control. **(E)** Cqd1 interacts homotypically. Mitochondria of yeast strains expressing Cqd1-3xMyc in the presence of untagged or 3xHA tagged Cqd1 were treated as described in (B). T, total lysate (5%); UB, unbound protein (5%); B, bound protein (100%). The asterisk indicates a cross reaction of the anti-Myc antibody. **(F)** Cqd1 interacts with other proteins than itself. Isolated mitochondria of a Cqd1-3xHA expressing strain were exposed to DMSO or the chemical crosslinkers DSG and MBS at the indicated concentrations. Samples were analyzed by immunoblotting with an anti-HA antibody. Arrows indicate Cqd1-containing crosslinks. **(G)** Cqd1 interacts with Om14. Mitochondria of wild type and a yeast strain expressing Om14-3xHA were analyzed as in (B). T, total lysate (2.5%); UB, unbound protein (2.5%); B, bound protein (100%). The asterisk indicates a cross reaction of the anti-Tom70 antibody. **(H)** Cqd1 interacts with Por1. Mitochondria of wild type or a yeast strain expressing Por1-3xHA were analyzed as in (B). T, total lysate (1%); UB, unbound protein (1%); B, bound protein (100%). The arrowhead indicates degradation products of Por1-3xHA. The asterisk indicates a cross reaction of the anti-Om45 antibody. **(I)** The contact site formed by Cqd1 and Om14 is independent of MICOS. Mitochondria of wild type, Om14-3×HA and Δ*mic60* Om14-3×HA strains were analyzed as in (B). T, total lysate (2.5%); UB, unbound protein (2.5%); B, bound protein (100%). The asterisk indicates a cross reaction of the anti-Mic60 antibody.

In order to analyze whether Cqd1 is a novel MICOS subunit we performed immunoprecipitation of Cqd1 and different MICOS subunits. Neither the isolation of Cqd1 resulted in co-isolation of the MICOS subunits Mic10 or Mic60 nor vice versa (Fig. 6B and C). Native gel electrophoresis revealed that Cqd1 is present in a high molecular weight complex of ca. 400 kDa, a size that is clearly different from the 1.5 MDa large MICOS complex (Fig. 6D). These results suggest that Cqd1 is part of a novel contact site rather than a new MICOS subunit.

To screen for possible interaction partners of Cqd1 we first tested whether it might interact homotypically. To this end a yeast strain was generated expressing simultaneously a 3xHA-tagged and a 3xMyc-tagged version of Cqd1. Immunoprecipitation of Cqd1-3xHA led to efficient co-isolation of Cqd1-3xMyc, revealing homotypic interactions of Cqd1 (Fig. 6E).

While the homotypic Cqd1 interaction could explain the formation of the high molecular weight complex (Fig. 6D), it does not explain the presence of Cqd1 in contact site fractions (Fig. 6A). Therefore, chemical crosslinking was used to get an indication of further interaction partners. While incubation with Di(N-succinimidyl) glutarate (DSG) did not reveal any crosslink partners, treatment with m-maleimidobenzoyl-N-hydoxysuccinimide ester (MBS) produced two prominent crosslinks (Fig. 6F).

These two crosslink products of about 80 and 100 kDa indicated interaction partners with a molecular weight of about 15 and 30 kDa. We identified Om14 and Por1, two highly abundant proteins forming a complex in the mitochondrial outer membrane (Lauffer et al., 2012; Sakaue et al., 2019), as promising candidates. Consistently, immunoprecipitation of Om14-3xHA allowed the co-isolation of Cqd1 and Por1 (Fig. 6G) and Cqd1 was co-isolated upon immunoprecipitation of Por1-3xHA (Fig. 6H). The coisolation efficiency of Cqd1 was even higher than the co-isolation efficiency of Om45, a previously identified interaction partner of Por1 in the outer membrane (Lauffer et al., 2012; Sakaue et al., 2019; Wenz et al., 2014). Of note, we were not able to co-purify Por1 by isolating Cqd1-3xHA (Fig. 6B) indicating that immunoprecipitation of Cqd1 impacts the integrity of the complex. Interestingly, Por1 was recently shown to form a contact site with the mitochondrial inner membrane protein Mdm31 (Miyata et al., 2018). Therefore, it might be that Cqd1 is a novel subunit of this complex. However, isolation Mdm31-3xHA did not result in co-isolation of Cqd1 (our unpublished observations).

Most of the so far identified contact sites between the mitochondrial inner and outer membranes depend on MICOS (Bohnert et al., 2012; Darshi et al., 2011; Harner et al., 2011; Hoppins et al., 2011; Korner et al., 2012; Modi et al., 2019; von der Malsburg et al., 2011; Xie et al., 2007; Zerbes et al., 2012). In particular, Mic60 is essential for the integrity of the MICOS complex as well as the formation of MICOS-dependent contact sites (Harner et al., 2011; Hoppins et al., 2011; von der Malsburg et al., 2011). Although we did not detect a physical interaction between Cqd1 and MICOS, the interaction of Cqd1 with Por1 and Om14 might still depend on the presence of an intact MICOS complex. To test this, we performed immunoprecipitation of Om14-3xHA expressed in the wild type background or the Δ*mic60* deletion background. We observed that immunoprecipitation of Om14-3xHA led to the successful co-isolation of Cqd1 and Por1 independent of the presence of Mic60 (Fig. 6I).

Taken together, we identified a novel contact site formed by the mitochondrial inner membrane protein Cqd1 and the previously described Por1-Om14 complex in the mitochondrial outer membrane (Lauffer et al., 2012; Sakaue et al., 2019). This new contact site exists independently of contact sites formed by MICOS.

### Overexpression of *CQD1* leads to altered mitochondrial architecture and morphology

Cells overexpressing *CQD1* from the strong *GAL* promoter showed a dramatically reduced growth rate (Fig. 7A). Western blot analysis using a Cqd1 specific antibody revealed the appearance of a slower migrating band, probably representing the accumulation of some Cqd1 precursor protein, upon overexpression (Fig. 7B, arrowhead). The levels of several other mitochondrial proteins involved in mitochondrial dynamics, protein import, mitochondrial architecture or respiration were not changed compared to wild type and the Δ*cqd1* deletion mutant (Fig. 7B). However, when we analyzed the presence of different mitochondrial protein complexes, we found that monomeric and dimeric F_1_F_0_ ATP synthase complexes were strongly reduced. In contrast, respiratory chain super complexes were not significantly changed (Fig. 7C).

**Figure 7.**
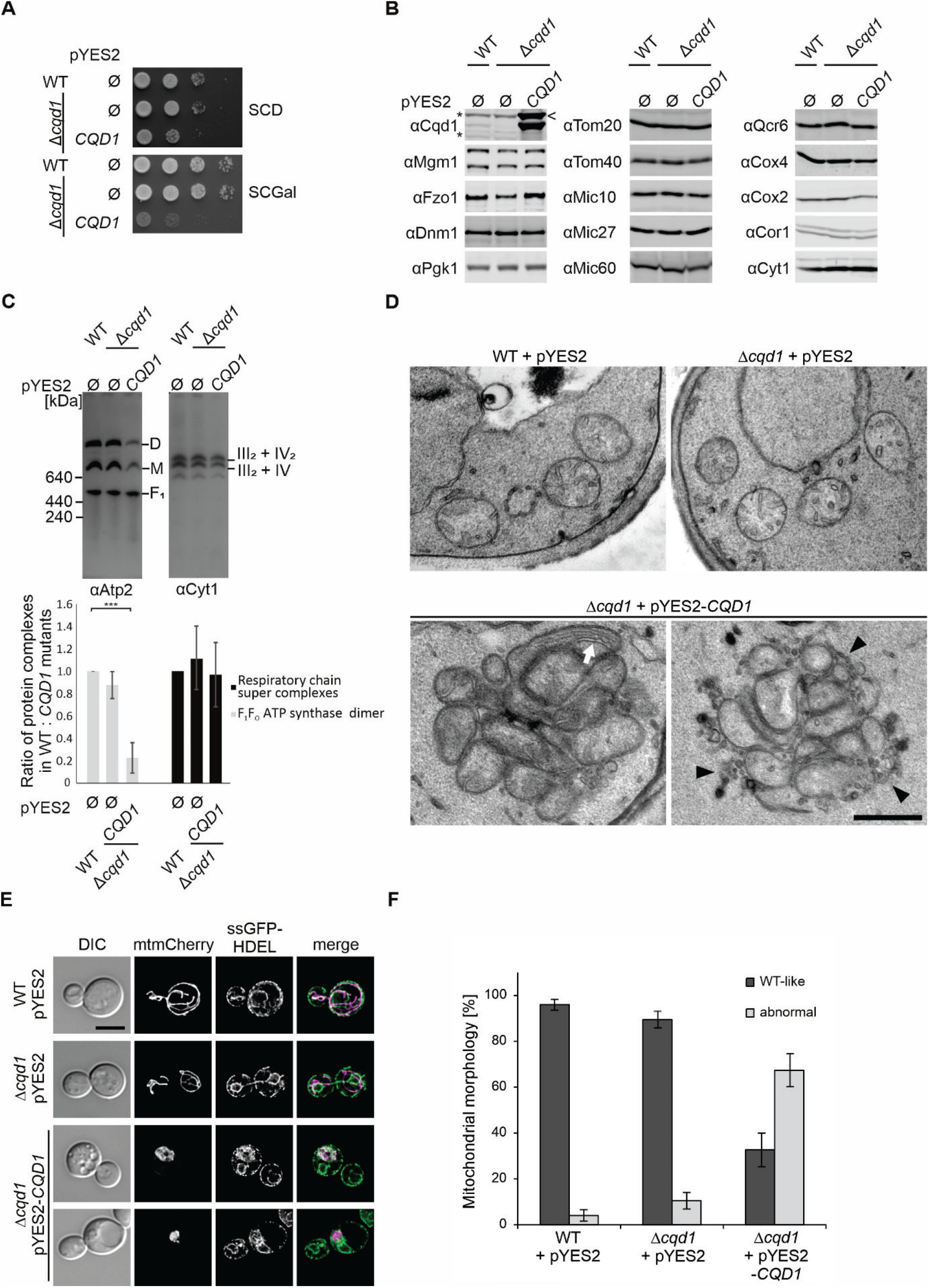
Overexpression of *CQD1* leads to dramatic changes in architecture and morphology of mitochondria. **(A)** Overexpression of *CQD1* is toxic. Wild type and Δ*cqd1* cells carrying an empty vector (pYES2) and Δ*cqd1* cells expressing *CQD1* from a high-copy plasmid with a galactose-inducible promoter (pYES2-*CQD1*) were grown in SCD and shifted to SCGal prior to analysis. **(B)** Deletion or overexpression of *CQD1* does not affect steady state levels of mitochondrial proteins. Whole cell extracts of wild type and yeast cells in which *CQD1* was deleted or overexpressed were analyzed by immunoblotting. Asterisks indicate cross reactions of the anti-Cqd1 antibody. The arrowhead indicates the potential precursor form of Cqd1. **(C)** Overexpression of Cqd1 results in strongly reduced levels of assembled F_1_F_O_ ATP synthase. Isolated mitochondria of cells grown in SCGal were lysed in buffer containing digitonin (3% w/v) and cleared lysates were subjected to BN-PAGE. The assembly of the F_1_F_O_ ATP synthase or the respiratory chain super complexes was analyzed by immunoblotting using antibodies against Atp2 or Cyt1. D, dimer of the F_1_F_O_ ATP synthase; M, monomer of the F_1_F_O_ ATP synthase; F_1_, F_1_ subcomplex of the F_1_F_O_ ATP synthase. (Upper panel) Immunodecorations of one representative experiment are shown. (Lower panel) Quantification of signal intensities of the F_1_F_O_ ATP synthase dimers and the respiratory chain super complexes (complex III dimer - complex IV monomer and complex III dimer - complex IV dimer) present in the indicated strains of three independent experiments were done with Image J software. Error bars indicate standard deviation. Asterisks represent p-values calculated by one-way ANOVA with subsequent Tukey’s multiple comparison test (***p≤0.001). **(D)** Overexpression of *CQD1* results in highly altered mitochondrial architecture. Cells were grown overnight in SCGal, subjected to chemical fixation with glutaraldehyde and osmium tetroxide, embedded in Epon, and ultrathin sections were analyzed by transmission electron microscopy. The white arrow highlights elongated inner membrane structures, black arrowheads highlight membranes presumably representing cross-sections of ER tubules. Scale bar, 500 nm. Additional electron micrographs are shown in Suppl. Fig. 2. **(E)** Overexpression of *CQD1* leads to the formation of mitochondria-ER clusters. Strains expressing mitochondria- targeted mCherry (mtmCherry) and ER-targeted GFP (ssGFP-HDEL) were grown to logarithmic growth phase in SCGal, fixed with formaldehyde, and examined by deconvolution fluorescence microscopy. Shown are maximum intensity projections of z stacks of entire cells (mitochondria) or of the center of the cells (ER, four consecutive z sections). DIC, *differential interference contrast.* Scale bar, 5 μm. **(F)** Strains expressing mitochondria-targeted GFP were analyzed as in (E) and mitochondrial morphology was quantified. Columns represent mean values from two independent experiments with three biological replicates per strain and at least 150 cells per replicate, error bars indicate standard deviation. Representative images are shown in Suppl. Fig. 3.

The F_1_F_O_ ATP synthase is essential for the establishment of mitochondrial inner membrane architecture (Klecker and Westermann, 2021). Absence of assembled, dimeric F_1_F_O_ ATP synthase results in the formation of septa that cross the matrix completely or onion-like structures (Davies et al., 2012; Habersetzer et al., 2013; Harner et al., 2016; Paumard et al., 2002; Rabl et al., 2009). Therefore, we asked whether the lack of assembled F_1_F_O_ ATP synthase complexes in the *CQD1* overexpression strain results in altered mitochondrial architecture. The Δ*cqd1* deletion strain appeared like wild type also in electron micrographs, whereas overexpression of *CQD1* led to massively distorted mitochondrial membrane architecture in many cells (Fig. 7D, Suppl. Fig. 2). Mitochondria were often aggregated into huge clusters and we frequently observed loss of normal cristae, in many cases accompanied by the formation of elongated inner membrane structures (Fig. 7D, arrow, Suppl. Fig. 2). Thus, aberrant mitochondrial architecture upon overexpression of Cqd1 complements the loss of assembled F_1_F_O_ ATP synthase.

Furthermore, in several cases we noticed membraneous structures within these clusters that might represent ER membranes (Fig. 7D, arrowheads, Suppl. Fig. 2). To test this, we simultaneously analyzed the morphology of mitochondria and the ER by fluorescence microscopy in wild type and the *CQD1* deletion and overexpression strains. When *CQD1* was overexpressed mitochondria in most cells did not appear as the characteristic tubular network that is spread all over the cell, but instead formed swollen clusters (Fig. 7E and F, Suppl. Fig. 3). Remarkably, these clusters indeed contained a large proportion of ER membranes (Fig. 7E).

In summary, overexpression of *CQD1* causes severe phenotypes on growth, assembly of the F_1_F_O_ ATP synthase, and mitochondrial architecture and morphology. Interestingly, the effect of overexpression of the inner membrane protein Cqd1 is apparently not only restricted to the inner membrane but also affects inter-organellar contacts.

## Discussion

In the present study, we characterized the highly conserved protein kinase-like domain containing protein Cqd1. All members of the UbiB family present in yeast, Cqd1, Cqd2 and Coq8, participate in coenzyme Q metabolism (Do et al., 2001; Johnson et al., 2005; Kemmerer et al., 2021; Reidenbach et al., 2018; Stefely et al., 2015; Subramanian et al., 2019; Xie et al., 2012). In contrast to the Δ*coq8,* mutant deletion of *CQD1* does not result in reduced production but affects distribution of coenzyme Q within the cellular membrane system (Kemmerer et al., 2021). The molecular role of Cqd1 in this process is unknown so far. However, membrane contacts would be ideally suited to facilitate the transition of these hydrophobic molecules (Kemmerer et al., 2021). The key finding of our study, the contact site formed by Cqd1, Por1 and Om14, is in good agreement with this hypothesis. Based on our results indicating that Cqd1 is also involved in phospholipid homeostasis, it is tempting to speculate that it might contribute to the import of phosphatidic acid, and possibly other lipids, into the inner membrane in addition to its role in coenzyme Q distribution.

We consider it unlikely that Cqd1 itself is a lipid transporter, as the absence of Cqd1 leads to increased export of coenzyme Q rather than block of transport (Kemmerer et al., 2021). However, it is conceivable that Cqd1 has an accessory or regulatory role and interacts with additional, yet unknown interaction partners in the inner membrane. This idea would be in line with models of other lipid transport systems, like the bacterial maintenance of lipid asymmetry (Mla) pathway first identified in *E. coli* (Malinverni and Silhavy, 2009). Here, the MlaFEDB complex is involved in the ATP-dependent transport of lipids between the bacterial inner and outer membranes (Malinverni and Silhavy, 2009; Thong et al., 2016). This complex is composed of the canonical proteins MlaE and MlaF and the auxiliary proteins MlaB and the hexameric MlaD in the inner membrane, and cooperates with MlaC, a soluble protein in the periplasm, and the outer membrane protein MlaA (Ekiert et al., 2017; Malinverni and Silhavy, 2009; Thong et al., 2016). Intriguingly, MlaA interacts with the porins OmpC and OmpF (Chong et al., 2015) and it was proposed that it works in both directions between the bacterial membranes (Ekiert et al., 2017; Malinverni and Silhavy, 2009). The Mla pathway represents a transport system based on transient protein interactions, in contrast to the here discovered permanent contact. However, it shares two striking features with Cqd1: an interaction with porin across two membranes and the regulation of bi-directional lipid transport.

In addition to the observed changes in phospholipid composition upon deletion of *CQD1,* overexpression of *CQD1* results in altered mitochondrial architecture. These changes are characterized by the formation of huge mitochondrial clusters and aberrant inner membrane structures. We and others observed similar alterations of inner membrane structure when analyzing yeast mutants lacking dimeric F_1_F_O_ ATP synthase (Davies et al., 2012; Harner et al., 2016; Paumard et al., 2002; Rabl et al., 2009). And indeed, overexpression of *CQD1* goes along with a strong reduction of assembled F_1_F_O_ ATP synthase. Furthermore, we observed that mitochondrial clusters intimately interact with ER membranes. This suggests that Cqd1 might have a function in mitochondrial architecture that goes beyond regulation of coenzyme Q and phosphatidic acid homeostasis. It remains a challenge for the future to reveal the molecular mechanisms in this process.

## Material and Methods

### Yeast strains and cell growth

*S. cerevisiae* YPH499 was used as wild type (WT). Chromosomal manipulations (knockouts, C-terminal 3xHA and 3xMyc tagging) were performed according to established procedures (Knop et al., 1999; Longtine et al., 1998). For the generation of deletion strains, the entire coding regions of the corresponding genes were replaced by the indicated marker cassettes. Double mutant strains were also generated by homologous recombination. The genotypes of the strains used in this study are listed in Supplementary Table 2.

For the generation of pRS316-*CQD1,* pRS316-*CQD1*-3xHA and pYES2-*CQD1* plasmids, the coding region of *CQD1* was amplified by PCR. The intron present in the *CQD1* open reading frame was removed when cloning the initial pYES2 construct. This construct served as template DNA for cloning of pRS316-*CQD1*. The coding region of *CQD1*-3xHA was amplified from yeast genomic DNA of the *CQD1*-3xHA strain. *CQD1*-3xHA was cloned under control of its endogenous promoter and the *ADH1* terminator into the pRS316 vector by enzymatic assembly of overlapping DNA fragments (Gibson, 2011). The point mutations K275A, D288A, and E330A in *CQD1* were introduced into pRS316-*CQD1* by site directed mutagenesis. For the generation of the pYX142-ssGFP-HDEL plasmid the *Sac*I/*Eco*RI fragment of pYX122-ssGFP-HDEL (Bockler and Westermann, 2014) was ligated into the *Sac*I/*Eco*RI sites of pYX142- mtGFP (Westermann and Neupert, 2000). The primers used in this study are listed in Supplementary Table 3.

Yeast cells were grown as indicated on YP medium supplemented with 2% glucose (YPD) or 3% glycerol (YPG) or synthetic medium supplemented with 2% glucose (SCD), 3% galactose (SCGal) or 3% glycerol (SCG) (Izawa and Unger, 2017).

Growth of the different strains was analyzed by the drop dilution assay. Cells were grown and kept in logarithmic growth phase, washed with water, diluted in water to an OD_600_ of 0.5 and serial dilutions were prepared (1:10; 1:100; 1:1,000). 5 μl of each dilution were spotted on agar plates with the indicated media. All strains without plasmids were grown initially in YPD liquid medium for 16 h. For analysis of cell growth on YP medium, cells were harvested after growth in YPD. When growth on synthetic medium was analyzed, cells were shifted from YPD to SCD 30 h prior to analysis. Cell carrying pYES2 plasmids were grown initially in SCD liquid medium for 16 h and shifted to SCGal for 7h before growth was analyzed.

### Whole cell extract

Whole cell extracts were prepared as described previously (Kushnirov, 2000). In brief, cells were grown in the indicated media to exponential growth phase (OD_600_ 0.5 - 1.0) and an equivalent of cells corresponding to 2.5 OD_600_ was collected by centrifugation. Harvested cells were resuspended in 200 μl of 0.1 M NaOH and incubated at room temperature for 5 min. Cell pellets were resuspended in 50 μl of SDS sample buffer and analyzed by SDS-PAGE and immunoblotting.

### Isolation of crude mitochondria

Cells were grown at 30°C in SCD, SCG, SCGal or YPG as indicated. Mitochondria were isolated by differential centrifugation as described previously (Izawa and Unger, 2017) with slight modifications. In brief, cells were harvested and washed with distilled water. Cells were treated for 10 min with DTT (10 mM final concentration) followed by treatment with zymolyase (200 U per gram wet weight of cells) for 30 min at 30°C with gentle agitation. Sphaeroplasts were opened by repeated pipetting in isotonic lysis buffer (20 mM MOPS-KOH pH 7.2, 1 mM EDTA, 0.6 M sorbitol, 0.2% (w/v) BSA, 1 mM phenylmethylsulfonyl fluoride (PMSF)). Mitochondria were harvested after a clarifying spin at 2,000 ×g and 4°C for 5 min by centrifugation at 14,000 ×g and 4°C for 10 min and resuspended in SM buffer (0.6 M sorbitol, 20 mM MOPS, pH 7.4).

### Sucrose gradient purification of mitochondria

Crude mitochondria were isolated as described above except that the final mitochondrial pellet was resuspended in 15 ml of buffer A (0.6 M Sorbitol, 50 mM MES pH 6.0) and re-isolated by centrifugation for 10 min at 17,000 xg and 4°C. Crude mitochondria were resuspended in 2 ml of buffer A and 10 mg were loaded on top of a sucrose step gradient (1.5 ml of 60%, 5 ml of 35%, 1.5 ml of 25%, and 1.5 ml of 15% sucrose in buffer A). Mitochondria were separated from other organelles by ultracentrifugation for 1 h at 134,000 xg and 4°C. Purified mitochondria were harvested from the boundary between 60% and 35% sucrose layers, resuspended in 15 ml of SM buffer and re-isolated by centrifugation for 15 min at 14,000 xg and 4°C. Mitochondria were resuspended in SM buffer to a final concentration of 10 mg/ml.

### Subfractionation of mitochondria

Vesicles consisting of pure mitochondrial outer membrane, mitochondrial inner membrane, and vesicles consisting of both membranes were generated and separated as described before (Harner, 2017) with slight modifications. 10 mg of freshly isolated crude mitochondria were resuspended in 1.6 ml of SM buffer. Mitochondria were swollen by addition of 16 ml swelling buffer (20 mM MOPS pH 7.4) and subsequent incubation for 30 min at 4°C under mild stirring. Sucrose concentration was adjusted to 0.55 M. Vesicles were generated by sonication (four times 30 seconds at 10% amplitude intermitted by 30 seconds breaks). After a clarifying spin for 20 min at 20,000 ×g and 4°C, vesicles were concentrated on the 2.5 M sucrose cushion by centrifugation at 118,000 ×g and 4°C for 100 min. Concentrated vesicles were harvested and the suspension was homogenized by dounce homogenization. Vesicles were separated by centrifugation for 12 hours at 200,000 ×g and 4°C through a sucrose step gradient (0.8 M, 0.96 M, 1.02 M, 1.13 M, 1.25 M sucrose in 20 mM MOPS-KOH pH 7.4 and 0.5 mM EDTA). The gradient was harvested and proteins were subjected twice to trichloroacetic acid precipitation (TCA; final concentration of 14.4%).

### Alkaline extraction

100 μg of isolated mitochondria were diluted in SM buffer to a concentration of 1 mg/ml. An equal volume of 200 mM sodium carbonate was added followed by an incubation on ice for 30 min. Membrane and soluble proteins were separated by centrifugation for 30 min at 91,000 ×g and 4°C. Proteins present in the supernatant were subjected to TCA precipitation. The membrane protein pellet and the precipitated soluble proteins were resuspended in SDS sample buffer and analyzed by SDS-PAGE and immunoblotting.

### Proteolytic susceptibility assay

100 μg of mitochondria were incubated in SM buffer (0.6 M sorbitol, 20 mM MOPS, pH 7.4), swelling buffer (20 mM MOPS, pH 7.4) or lysis buffer (1% (v/v) Triton X-100, 20 mM MOPS, pH 7.4) for 20 min on ice. Proteinase K was added (final concentration of 0.2 mg/ml) as indicated and samples were incubated on ice for 15 min. Proteolysis was stopped by addition of PMSF to a final concentration of 4 mM. Samples were centrifuged at 17,000 ×g and 4°C for 20 min. Pellets were resuspended in SM buffer and subjected to TCA precipitation. Samples were analyzed by SDS-PAGE and immunoblotting.

### Blue native gel electrophoresis (BN-PAGE)

150 μg (MICOS complex) or 50 μg (all other mitochondrial protein complexes) of isolated mitochondria were resuspended in 15 μl (MICOS complex) or 50 μl (all other mitochondrial protein complexes) of BN buffer (50 mM NaCl, 50 mM imidazole-HCl, 2 mM 6-aminohexanoic acid, 1 mM EDTA, 1 mM PMSF, pH 7.0) (Wittig et al., 2006) supplemented with 3% digitonin. After a clarifying spin for 15 min at 14,000 ×g and 4°C 2 μl (MICOS complex) or 7 μl (all other mitochondrial protein complexes) G-250 Sample Additive (Invitrogen, Waltham, Massachusetts, USA) containing 40% glycerol were added. Protein complexes were separated on 3-12% NativePAGE Novex Bis-Tris gels (Invitrogen). Electrophoresis was performed for 30 min at 150 V followed by 2 hours at 250 V. Proteins were transferred to PVDF membrane by wet blotting for 2 hours at 30 mV.

### Chemical crosslinking

Isolated mitochondria were diluted in 100 μl of SI buffer (50 mM HEPES-KOH, 0.6 M sorbitol, 75 mM KCl, 10 mM Mg(Ac)_2_, 2 mM KH_2_PO_4_, 2.5 mM EDTA, 2.5 mM MnCl_2_, pH 7.2) to a concentration of 1 mg/ml. Disuccinimidyl glutarate (DSG) and m-maleimidobenzoyl-N-hydroxysuccinimide ester (MBS) were added from freshly prepared stock solutions to the final concentration of 200 or 400 μM and incubated on ice for 30 min. The reaction was stopped by the addition of glycine to a final concentration of 100 mM followed by incubation on ice for 10 min. Mitochondria were re-isolated by centrifugation at 17,000 xg and 4°C for 10 min. The pellets were resuspended in SDS sample buffer and analyzed by SDS-PAGE and immunoblotting.

### Immunoprecipitation assay

3 mg mitochondria (except for Cqd1-3×HA, 1 mg of mitochondria) were lysed in IP buffer (50 mM Tris-HCl pH 7.4, 50 mM NaCl) containing 1% (w/v) digitonin and 1 mM PMSF. After a clarifying spin for 10 min at 12,000 xg and 4°C, lysates were incubated with anti-HA agarose beads (20 μl of beads per 1 mg of proteins) (Merck, Darmstadt, Germany) for 2 h. Anti-HA agarose beads were washed 3 times with 500 μl IP buffer containing 0.1% (w/v) digitonin and 1 mM PMSF before and after incubation with lysates. The indicated amount of total lysate and unbound material was taken, subjected to TCA precipitation and resuspended in SDS sample buffer. Bound proteins were eluted with SDS sample buffer at 95°C for 10 min. Fractions were analyzed by SDS-PAGE followed by immunoblotting.

### Fluorescence microscopy

For visualizing mitochondria, the mitochondrial presequence of *Neurospora crassa* subunit 9 of the F_1_F_O_ ATP synthase was fused to mKate2. The nucleotide sequence was inserted into the HO locus of the yeast genome and expressed constitutively under the control of the *PGK1* promoter. Yeast cells were grown overnight in YPD at 30°C. The next day, cells were diluted in synthetic media containing 2% glucose and were kept in logarithmic phase for 24 hours. Cells corresponding to an OD_600_ of 1 were harvested by centrifugation at 2,500 xg for 3 min. Cells were vortexed for 1 min and washed with 1 ml of sterile 1x PBS. Cell pellets were resuspended in 200 μl 1x PBS and immobilized on μ-slides (Ibidi, Gräfelfing, Germany) coated with 1 mg/ml of concanavalin A. After immobilizing, cells were covered with 400 μl SCD medium. Microscopy was performed at 30°C on a Nikon Ti2-Eclipse microscope (Nikon, Tokyo, Japan) equipped with a CFI Apochromat TIRF 100x/1.49 NA oil objective and a TwinCam LS dual camera splitter attached to two Photometrics Prime 95B 25 mm cameras (Teledyne Photometrics, Tucson, USA).

For visualization of mitochondria or mitochondria and the ER upon *CQD1* overexpression, strains carried either pYX142-mtGFP (Westermann and Neupert, 2000) or pYX232-mtmCherry (Dirk Scholz, University of Bayreuth) and pYX142-ssGFP-HDEL, respectively. Cells were grown overnight in galactose-containing synthetic complete medium supplemented with 0.1% glucose, shifted to galactose-containing synthetic complete medium without glucose, incubated until logarithmic growth phase and fixed with 3.7% formaldehyde. Microscopy was performed using a Leica DMi8 fluorescence microscope (Leica Microsystems GmbH, Wetzlar, Germany) equipped with a HC PL APO 100x/1.40 oil objective, a Lumencor SPECTRA X light source and fluorescence filter sets (TXR Cube ex. 540-580 nm; em. 592-668 nm and FITC Cube ex. 460-500 nm; em. 512-542 nm). Images were taken with a Leica DFC9000 GT VSC-07400 sCMOS camera. For microscope settings, image generation, and processing (cropping, maximum intensity projection), the Leica LAS X software (version 3.6.0.20104, Leica Microsystems GmbH, Wetzlar, Germany) was used. For deconvolution of z stacks the Deconvolution Software Huygens Essential (version 18.10, Scientific Volume Imaging, Hilversum, The Netherlands) was used. For adjustment of brightness and contrast and the overlay of different channels Adobe Photoshop CS6 (Adobe Systems) was used. For simultaneous analysis of the morphology of mitochondria and the ER, z stacks were recorded with a z-step size of 213 nm. In the case of mitochondria, maximum intensity projections of z stacks of entire cells are shown, whereas for the ER maximum intensity projections of four consecutive z sections of the center of the cells are shown.

### Electron microscopy

For electron microscopy, cells were first grown in synthetic complete medium containing glucose as carbon source and then shifted to synthetic complete medium containing galactose as carbon source, in which they were grown overnight. Chemical fixation of yeast cells with glutaraldehyde and osmium tetroxide, dehydration, Epon embedding, and subsequent steps of sample preparation for electron microscopy were performed as described previously (Unger et al., 2017). Electron micrographs were taken with a JEOL JEM-1400 Plus transmission electron microscope operated at 80 kV, a JEOL Ruby CCD camera (3296 x 2472 pixels), and the TEM Center software Ver.1.7.12.1984 or Ver.1.7.19.2439 (JEOL, Tokyo, Japan).

### Lipid analysis

Lipidomics analyses were performed as described in Papagiannidis et al. (2021). Aliquots corresponding to 1500-2000 pmol total lipid were subjected to acidic Bligh and Dyer extractions performed in the presence of internal lipid standards from a master mix containing 40 pmol d7-PC mix (15:0/18:1-d7, Avanti Polar Lipids), 25 pmol PI (17:0/20:4, Avanti Polar Lipids), 25 pmol PE and 15 pmol PS (14:1/14:1, 20:1/20:1, 22:1/22:1, semi-synthesized as described in Özbalci et al, 2013), 20 pmol PA (PA 17:0/20:4, Avanti Polar Lipids) 5 pmol PG (14:1/14:1, 20:1/20:1, 22:1/22:1, semi-synthesized as described in Ozbalci et al. (2013), 25 pmol CL 57:4 (14:1/14:1/14:1/15:1, Avanti Polar Lipids) and 25 pmol MLCL (16:0/16:0/16:0, Echelon). Lipids recovered in the organic extraction phase were evaporated by a gentle stream of nitrogen. Prior to measurements, lipid extracts were dissolved in 10 mM ammonium acetate in methanol, diluted 1:10 and transferred into Eppendorf twin.tec 96-well plates. Mass spectrometric measurements were performed in positive ion mode on an AB SCIEX QTRAP 6500+ mass spectrometer equipped with chip-based (HD-D ESI Chip, Advion Biosciences) nanoelectrospray infusion and ionization (Triversa Nanomate, Advion Biosciences) as described (Özbalci et al, 2013). The following precursor ion scanning (PREC) and neutral loss scanning (NL) modes were used for the measurement of the various lipid classes: +PREC 184 (PC), +NL141 (PE), +NL185 (PS), +NL277 (PI), +NL189 (PG), +NL115 (PA). Mass spectrometry settings: Resolution: unit, low mass configuration; data accumulation: 400 MCA; curtain gas: 20; interface heater temperature: 60°C; CAD: medium. Data evaluation was done using LipidView (Sciex) and ShinyLipids, a software developed in house. For MS analysis of CL and MLCL, dried lipids were re-dissolved in 40% UPLC solvent B (90% 2-propanol/10% acetonitrile/0.1% formic acid/10 mM NH4HCO3) and transferred to silanized glass inserts (Phenomenex) using Hamilton syringes. The glass inserts were placed in Eppendorf tubes and centrifuged in an Eppendorf centrifuge at 3000 rpm for 3 min. Lipid samples were then subjected to UPLC-ESI-MS/MS analysis performed on an Ultimate® 3000 LC system (Dionex, Thermo Fisher Scientific) coupled to a QExactive Hybrid Quadrupole-Orbitrap instrument (Thermo Scientific). For LC separations, an ACQUITY UPLC CSH C18 1.7μm, 1.0 x 150 mm column (Waters) was used. The column oven temperature was set to 55°C, the temperature of the autosampler was set to 20°C. The flow rate used was 100 μl/min. The solvent composition used was as follows: 60% acetonitrile/40% H_2_O/0.1% formic acid/10 mM NH_4_HCO_3_ (solvent A), 90% 2-propanol/10% acetonitrile/0.1% formic acid/10 mM NH_4_HCO_2_ (solvent B). The starting solvent composition was 50% solvent B/50% solvent A. The conditions of the gradient were as follows: 7 min: 90% solvent B, 7-17 min: 90% solvent B, 17.1 min 50% solvent B and 17.1-25 min: 50% solvent B. The MS analyses were performed in the positive ion mode. CL and MLCL species were measured as [M+NH_4_^+^] ions. The following ESI source parameters were used: sheath gas flow rate: 4 (a.u.), auxiliary gas flow rate: 0, sweep gas flow rate: 0, spray voltage: 1.5 kV, capillary temperature: 200°C, S-lens RF level: 50. Full-MS scans were recorded using the following parameters: resolution: 140,000 (at m/z 200) AGC-target: 1e6, maximum IT: 200 ms, scan range: m/z 500-2000. Data evaluation was performed using MassMap.

### Statistics

Statistics were calculated using the software GraphPad Prism (version 5; GraphPad, San Diego, CA, USA). Mass spectrometry analysis was performed in quadruplicate. Data are represented as mean ± standard deviation. For statistical analysis, data was first tested for normal distribution using Shapiro-Wilk normality test. P-Values were calculated using Mann Whitney test. Analysis of Mdj1 precursor accumulation and Mgm1 processing in the various yeast strains was performed in triplicate. Data are represented as mean ± standard deviation. P-Values were calculated using an unpaired Student’s t test. Analysis of the steady state level of Cqd1 variants expressed from pRS316 under the control of the endogenous Cqd promotor as well as the level of assembled mitochondrial protein complexes were performed in triplicates. Data are represented as mean ± standard deviation. P-Values were calculated using an one-way ANOVA test with subsequent Tukey’s multiple comparison test. *p* < 0.05 was considered statistically significant.

## Supporting information

Khosravi at al. Supplemental Information

## Acknowledgements

We thank Rita Grotjahn (University of Bayreuth) for help with electron microscopy. M.E.H. thanks Dr. Christof Osman, Ludwig-Maximilians University, Munich, for providing his microscope and support. M.E.H. is greatly thankful for the helpful discussion with Dr. Bill Wickner, Dartmouth Medical School, Hanover. Furthermore, M.E.H. thanks Dr. Michael Kiebler, Ludwig-Maximilians, University, Munich, for generous and extensive support.

## Author contributions

S.K., X.C., A.K.U, J.F., T.K., M.E.H., conception and design, acquisition of data, analysis and interpretation of data, drafting or revising the article; T.S., C.L., R.S., acquisition of data, analysis and interpretation of data, revising the article; B.B., B.W., W.N., conception and design, analysis and interpretation of data, drafting or revising the article.

