## Supplementary material for "The UbiB family member Cqd1 forms a novel membrane contact site in mitochondria": Khosravi at al. Supplemental Information

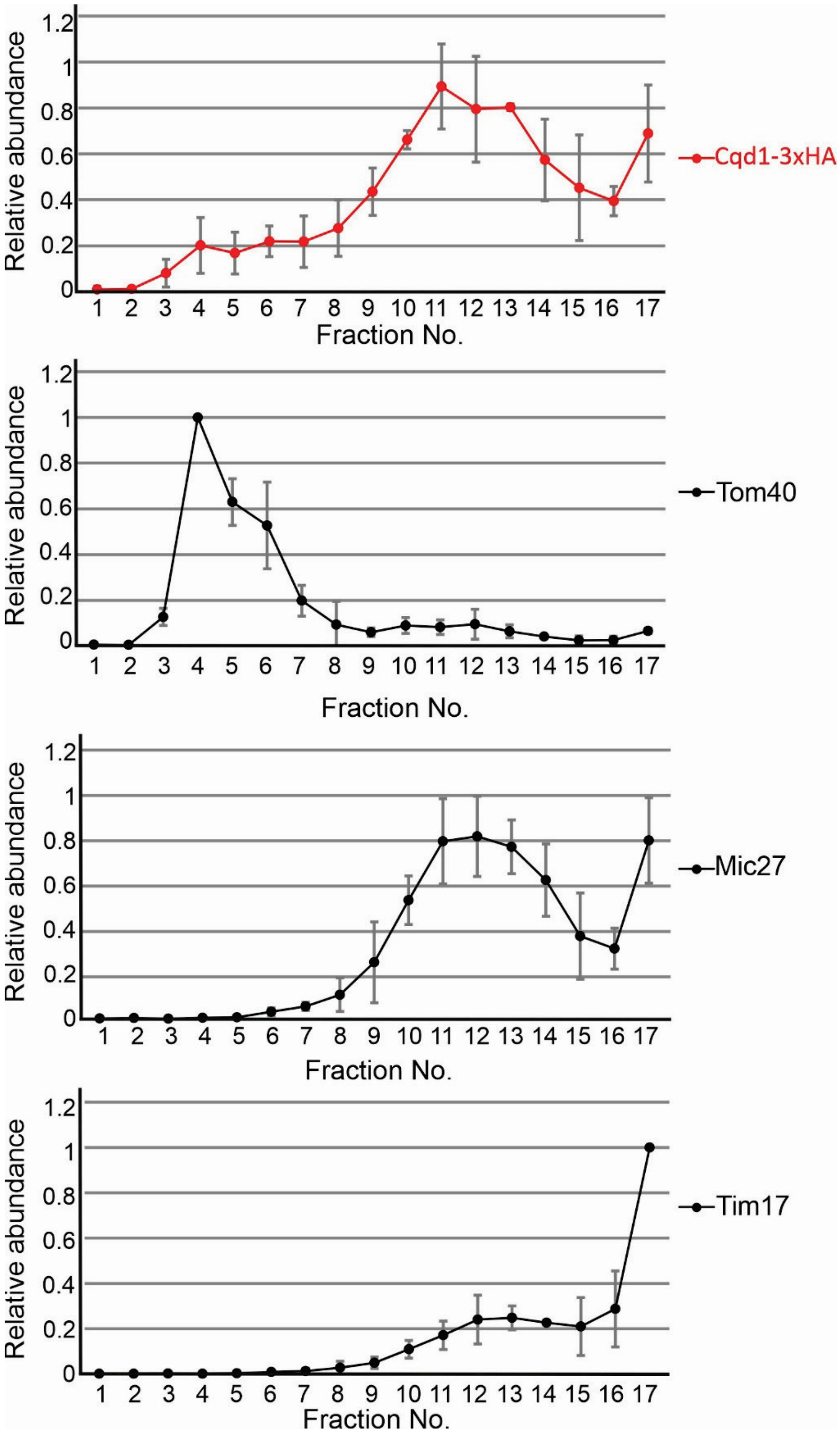


**Supplementary Figure 1. Cqd1 is enriched in fractions containing contact site proteins.**

The average of relative protein abundance in three independent experiments described in Fig. 6A is shown. The graph shows mean values of the distribution of Cqd1-3xHA and the marker proteins for the OM (Tom40), the IM (Tim17) and contact sites (Mic27). Error bars indicate standard deviation.

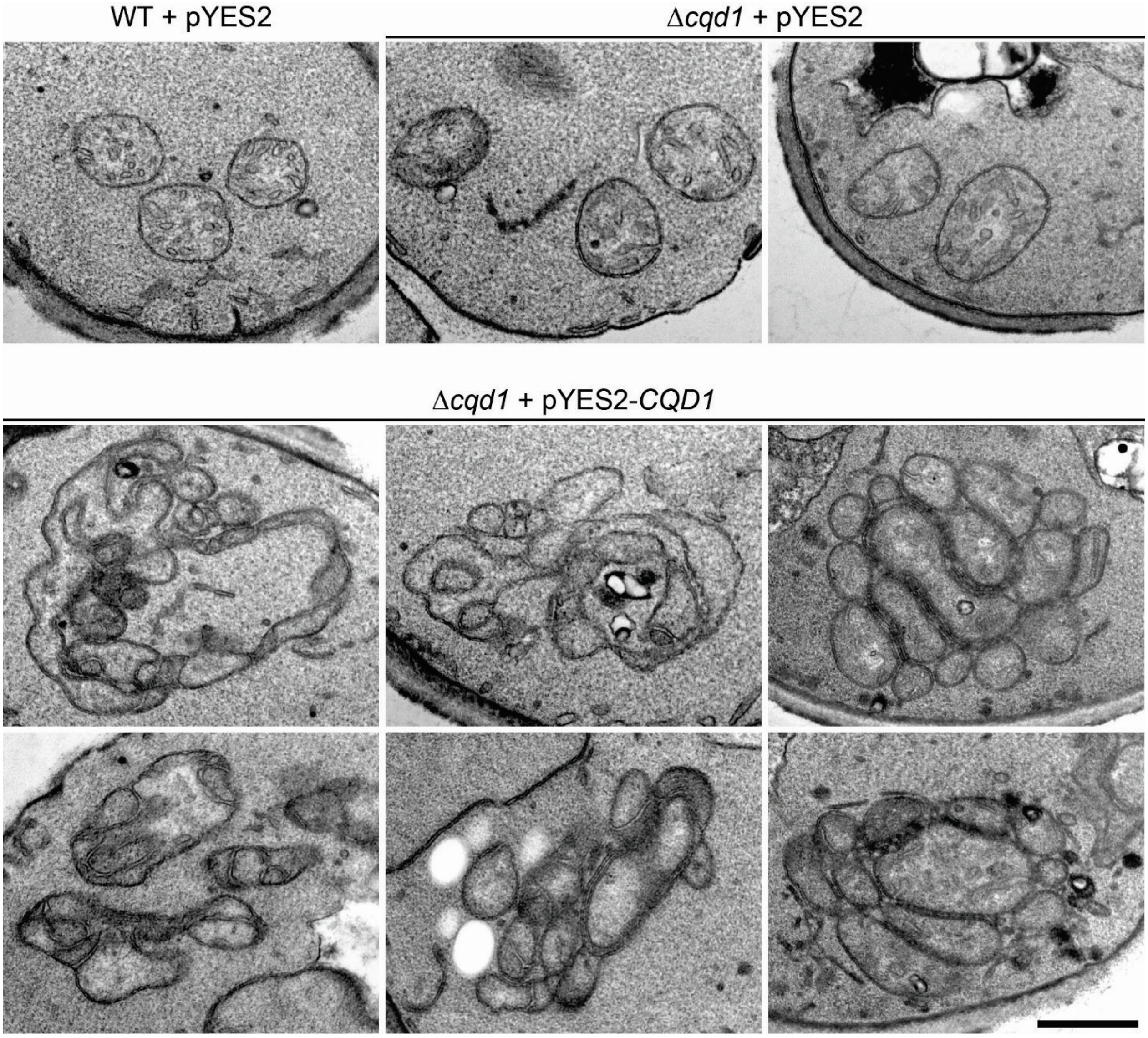


**Supplementary Figure 2. Cqd1-overexpression is associated with altered mitochondrial architecture.**

WT and Δ*cqd1* cells carrying the indicated plasmids were grown and analyzed by electron microscopy as described in Fig. 7D. Scale bar, 500 nm.

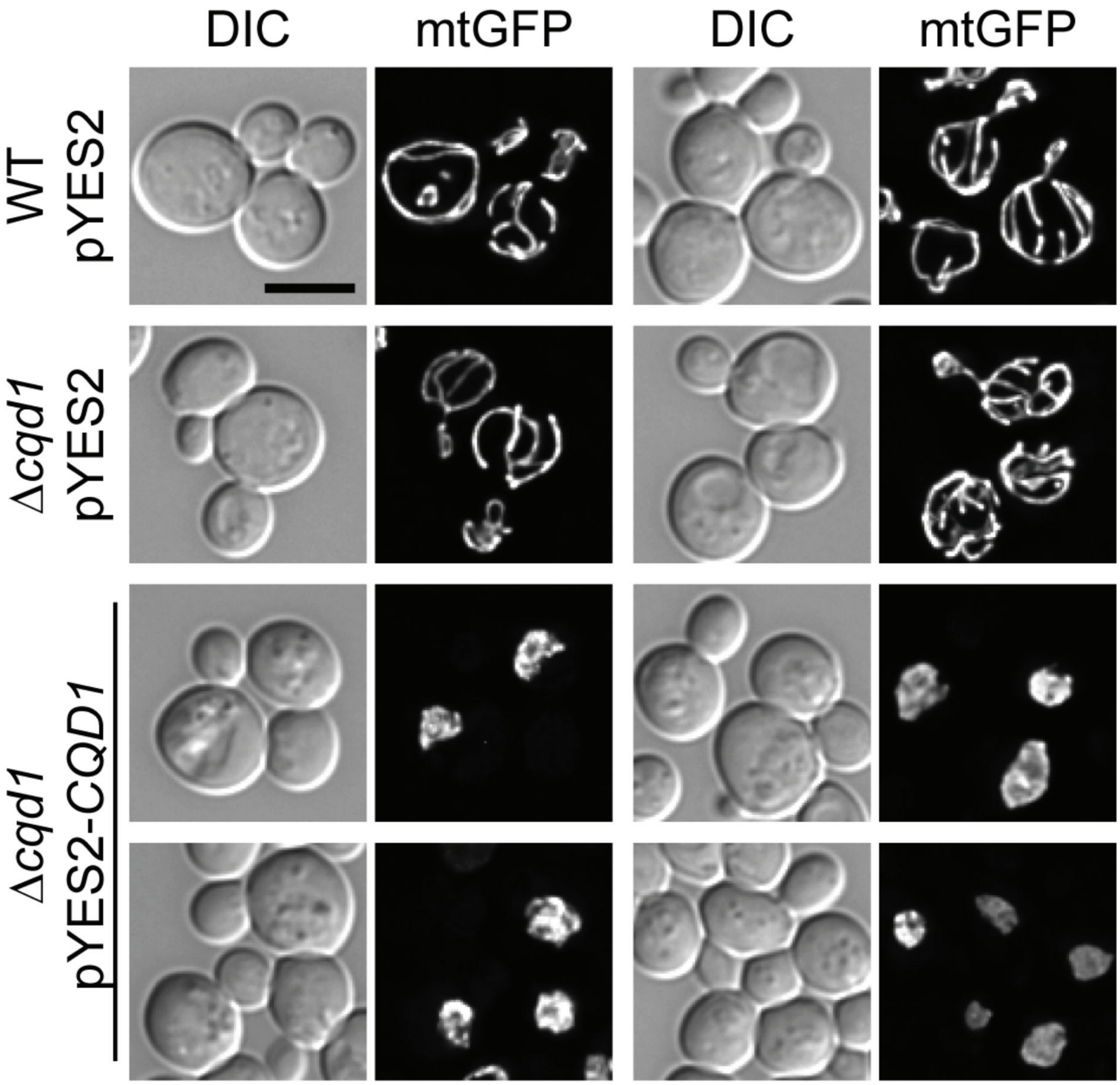


**Supplementary Figure 3. Overexpression of Cqd1 causes changes in mitochondrial morphology.**

WT and Δ*cqd1* cells carrying the indicated plasmids and expressing mitochondria-targeted GFP (mtGFP) were grown and analyzed as described in Fig. 7E. Shown are DIC (differential interference contrast) images and maximum intensity projections of deconvolved z stacks (mtGFP) from representative cells. Scale bar, 5 µm.

**Supplementary Table 1. Relative abundance of mitochondrial phospholipids.**

Values of the graph shown in Figure 3. PC, phosphatidylcholine; PE, phosphatidylethanolamine; PS, phosphatidylserine; PI, phosphatidylinositol; CL, cardiolipin; PG, phosphatidylglycerol; PA, phosphatidic acid; MLCL, monolysocardiolipin.

| average mol % lipid | | | | | | | | |
|---|---|---|---|---|---|---|---|---|
| **Sample** | **PC** | **PE** | **PS** | **PI** | **PG** | **PA** | **CL** | **MLCL** |
| WT | 36.17 | 27.30 | 5.05 | 16.48 | 0.15 | 4.17 | 10.45 | 0.22 |
| Δ*ups1* | 39.34 | 30.59 | 5.18 | 15.55 | 0.13 | 3.03 | 6.15 | 0.04 |
| Δ*cqd1* | 40.28 | 25.68 | 4.85 | 15.01 | 0.18 | 3.19 | 10.65 | 0.16 |
| Δ*ups1* Δ*cqd1* | 41.17 | 27.69 | 6.94 | 14.12 | 0.12 | 3.41 | 6.49 | 0.07 |
| **Standard deviation** | | | | | | | | |
| **Sample** | **PC** | **PE** | **PS** | **PI** | **PG** | **PA** | **CL** | **MLCL** |
| WT | 5.29 | 3.20 | 0.66 | 4.15 | 0.04 | 0.48 | 1.41 | 0.11 |
| Δ*ups1* | 3.12 | 3.80 | 1.22 | 1.81 | 0.10 | 0.16 | 0.61 | 0.03 |
| Δ*cqd1* | 1.50 | 2.30 | 0.62 | 1.49 | 0.06 | 0.46 | 1.04 | 0.04 |
| Δ*ups1* Δ*cqd1* | 2.36 | 2.17 | 2.15 | 2.27 | 0.04 | 0.80 | 2.13 | 0.07 |

**Supplementary Table 2. *S. cerevisiae* strains used in this study.**

| Strain name | Genotype | Reference |
|---|---|---|
| YPH499 | *MAT*a *ade2-101 his3-Δ200 leu2- trp1-Δ63 ura3-52 lys2-801* | (Sikorski and Hieter, 1989) |
| Δ*cqd1* | YPH499 *cqd1Δ::HIS3MX6* | This study |
| Δ*cqd2* | YPH499 *cqd2Δ::HIS3MX6* | This study |
| Δ*ups1* | YPH499 *ups1Δ::kanMX4* | This study |
| Δ*ups1* | YPH499 *ups1Δ::LEU2 (K. lactis)* | This study |
| Δ*tam41* | YPH499 *tam41Δ::LEU2 (K. lactis)* | This study |
| Δ*pgs1* | YPH499 *pgs1Δ::LEU2 (K. lactis)* | This study |
| Δ*gep4* | YPH499 *gep4Δ::LEU2 (K. lactis)* | This study |
| Δ*crd1* | YPH499 *crdΔ::LEU2 (K. lactis)* | This study |
| Δ*taz1* | YPH499 *taz1Δ::LEU2 (K. lactis)* | This study |
| Δ*cld1* | YPH499 *cld1Δ::LEU2 (K. lactis)* | This study |
| Δ*ups2* | YPH499 *ups2Δ::LEU2 (K. lactis)* | This study |
| Δ*mdm35* | YPH499 *mdm35Δ::LEU2 (K. lactis)* | This study |
| Δ*psd1* | YPH499 *psd1Δ::LEU2 (K. lactis)* | This study |
| Δ*ups1* Δ*cqd1* | YPH499 *cqd1Δ::HIS3MX6 ups1Δ::LEU2 (K. lactis)* | This study |

| | | |
|---|---|---|
| *Δups1 Δcqd1* | YPH499 *cqd1Δ::HIS3MX6 ups1Δ::kanMX4* | This study |
| *Δtam41 Δcqd1* | YPH499 *cqd1Δ::HIS3MX6 tam41Δ::LEU2 (K. lactis)* | This study |
| *Δpgs1 Δcqd1* | YPH499 *cqd1Δ::HIS3MX6 pgs1Δ::LEU2 (K. lactis)* | This study |
| *Δgep4 Δcqd1* | YPH499 *cqd1Δ::HIS3MX6 gep4Δ::LEU2 (K. lactis)* | This study |
| *Δcrd1 Δcqd1* | YPH499 *cqd1Δ::HIS3MX6 crd1Δ::LEU2 (K. lactis)* | This study |
| *Δtaz1 Δcqd1* | YPH499 *cqd1Δ::HIS3MX6 taz1Δ::LEU2 (K. lactis)* | This study |
| *Δcld1 Δcqd1* | YPH499 *cqd1Δ::HIS3MX6 cld1Δ::LEU2 (K. lactis)* | This study |
| *Δups2 Δcqd1* | YPH499 *cqd1Δ::HIS3MX6 ups2Δ::LEU2 (K. lactis)* | This study |
| *Δmdm35 Δcqd1* | YPH499 *cqd1Δ::HIS3MX6 mdm35Δ::LEU2 (K. lactis)* | This study |
| *Δpsd1 Δcqd1* | YPH499 *cqd1Δ::HIS3MX6 psd1Δ::LEU2 (K. lactis)* | This study |
| *Δcqd1 Δcqd2* | YPH499 *cqd2Δ::HIS3MX6 cqd1Δ::kanMX4* | This study |
| *Δups1 Δcqd2* | YPH499 *ups1Δ::kanMX6 cqd2Δ::hphNT1* | This study |
| *Δups1 Δcqd2 Δcqd1* | YPH499 *cqd2Δ::HIS3MX6 cqd1Δ::kanMX4 ups1Δ::LEU2 (K. lactis)* | This study |
| *CQD1-3xHA* | YPH499 *CQD1-3xHA::HIS3MX6* | This study |
| *CQD1-3xMyc* | YPH499 *CQD1::3xMyc-TRP1 (K. lactis)* | This study |
| *POR1-3xHA* | YPH499 *POR1-3xHA::HIS3MX6* | This study |
| *OM14-3xHA* | YPH499 *OM14-3xHA::HIS3MX6* | This study |
| *Δmic60 OM14-3xHA* | YPH499 *mic60Δ::HIS3MX6 OM14-3xHA::TRP1 (K. lactis)* | This study |
| WT mKate | YPH499 *ho-PGK1pr-su9-KATE2::kanMX4* | This study |
| *Δcqd1* mKate | YPH499 *cqd1Δ::HIS3MX6 ho-PGK1pr-su9-KATE2::kanMX4* | This study |
| *Δups1* mKate | YPH499 *ups1Δ::LEU2 (K. lactis) ho-PGK1pr-su9-KATE2::kanMX4* | This study |
| *Δups1 Δcqd1* mKate | YPH499 *ups1Δ::LEU2 (K. lactis) cqd1Δ::HIS3MX6 ho-PGK1pr-su9-KATE2::kanMX4* | This study |
| *Δups1CQD1-3xHA* | YPH499 *cqd1Δ::CQD1-3xHA::HIS3MX6 ups1::LEU2 (K. lactis)* | This study |
| *MIC10-3xHA* | YPH499 *MIC10-3xHA::HIS3MX6* | (Harner et al., 2011) |
| *MIC60-3xHA* | YPH499 *MIC60-3xHA::kanMX4* | (Harner et al., 2011) |

**Supplementary Table 3. Primers used in this study.**

| Construct | Primer name | Sequence |
|---|---|---|
| pYES2-*CQD1* | *CQD1*(SacI)_for | CCCGAGCTCATGTCATTTTTAAAGTTCGC |
| | *CQD1*-int(EcoRI)_rev | CCCGAATTCACGTAATATAGGCAGCGAACTTTCAAATAAGTCCAAG |
| | *CQD1*-int(EcoRI)_for | CGTGAATTCGGGTTTAAGC |
| | *CQD1*(NotI)_rev | CCCGCGGCCGCTTAATAATTAGGACACAATTG |
| pRS316-*CQD1* | F HindIII *CQD1* (gib) | GCAGGAATTCGATATCAAGCTTGGTACTGGAAAGATCGCGTTC |
| | R XhoI *CQD1* | CTTACCGGGCCCCCCCTCGACGTACCGTTGCCTTATTGTTC |
| pRS316-*cqd1(K275A)* | Cqd1_K275A_for | GCGATCTTGCATCCAAATGTAAG |
| | Cqd1_K275A_rev | GATGGCACACCAACGATTTC |
| pRS316- *cqd1(D288A)* | Cqd1_D288A_for | GCTTTGAAAATAATGAAATTCTG |
| | Cqd1_D288A_rev | TCTCCGGATCTGAGATCTTAC |
| pRS316- *cqd1(E330A)* | Cqd1_E330A_for | GCGGCGTTAAACCTGGAAAG |
| | Cqd1_E330A_rev | AATTCTTAGATCCAACTGAATA |
| pRS316-*CQD1*-3xHA | F HindIII *CQD1* (gib) | GCAGGAATTCGATATCAAGCTTGGTACTGGAAAGATCGCGTTC |
| | Gib(XhoI)*ADH1*terR1 | ACCGGGCCCCCCCTCGAGGTAGAGGTGTGGTCAATAAG |
